# Spatiotemporal proteomics reveals dynamic antagonistic gradients shaping signalling waves

**DOI:** 10.1101/2025.09.05.674076

**Authors:** Wilke H. M. Meijer, Virginia Andrade, Suzan Stelloo, Wouter M. Thomas, Marek J. van Oostrom, Eveline F. Ilcken, Kim T. J. Peters, Michiel Vermeulen, Katharina F. Sonnen

## Abstract

Embryonic development is driven by dynamic protein networks, yet how these dynamics shape morphogenesis remains incompletely understood. Somitogenesis, the rhythmic segmentation of vertebrate embryos, is governed by signalling gradients and oscillations in the presomitic mesoderm (PSM)^1,2^, but the corresponding protein dynamics are largely unknown. Perturbations in this process cause congenital spine disorders and can result in embryonic lethality^3^. Here, we introduce an integrated proteomics and microfluidics approach to resolve spatiotemporal protein expression in the developing mouse tail. To this end, we established a microfluidic system to synchronize oscillations in embryo tails grown in 3D, which we combined with mass-spectrometry and RNA sequencing. We uncover novel oscillatory proteins and differentially expressed genes along the anteroposterior axis. Building on this dataset, we identify a previously unrecognized antagonistic, dynamic ligand–receptor expression pattern in R-Spondin/LGR signalling explaining how Wnt-oscillation amplitude increases despite decreasing ligand levels in anterior PSM. Dynamic ligand expression was validated in mouse gastruloids. Perturbation of ligand dynamics reduced oscillation amplitude and impaired somite formation. Our study reveals a novel regulatory strategy in which dynamic antagonistic gradients fine-tune signalling strength, providing new mechanistic insight into how protein dynamics control tissue patterning. We anticipate our dataset to serve as foundation for mechanistic investigations of mammalian somitogenesis including the role of mechanics and metabolism. More broadly, our approach combining microfluidics-based synchronization of signalling in multicellular systems with omics analyses can be applied to study dynamics in other contexts such as in tissue homeostasis^4^ and regeneration^5^.

## Main

Embryonic development is a highly dynamic process, in which networks of proteins interplay to orchestrate cell fate decisions and morphogenesis with spatiotemporal precision. Somitogenesis in vertebrate embryos is the sequential segmentation of the presomitic mesoderm (PSM) into somites, blocks of tissue giving rise to vertebrae, muscle and skin. Perturbations in somitogenesis disrupt vertebral segmentation, leading to congenital spine defects and, in severe cases, embryonic lethality^3,6^. It is regulated by morphogen gradients along the anteroposterior (AP) axis of the PSM and signalling oscillations, collectively termed the segmentation clock^1,7,8^. According to the clock and wavefront model^9^, oscillations and gradients determine the timing and spacing of segmentation, respectively. Oscillations between neighbouring cells are slightly phase-shifted, resulting in kinematic waves travelling from posterior to anterior PSM, where a new pair of somites is formed every 2.5 hours in mice and 5-7 hours in human^10–13^.

Several signalling pathways regulating somitogenesis have been identified using in situ hybridization, transcriptomics and live imaging approaches^7,8,10,14–19^. These studies defined the segmentation clock as oscillatory Wnt, Notch and FGF signalling, while opposing gradients of Wnt/FGF (high posterior) and retinoic acid (high anterior) were also described^7,14,17,20–22^ (Fig. 1A, left panel). Together, these findings support a model in which somite formation in anterior PSM depends on the interaction of signalling gradients with periodic activity of the segmentation clock to induce differentiation and mesenchymal-to-epithelial transition^3,19,23–25^. Importantly, the Wnt–Notch phase relationship shifts from out-of-phase posteriorly to in-phase anteriorly, and microfluidic experiments demonstrated that this dynamic coupling is essential for proper somite formation^14,18^.

**Figure 1.**
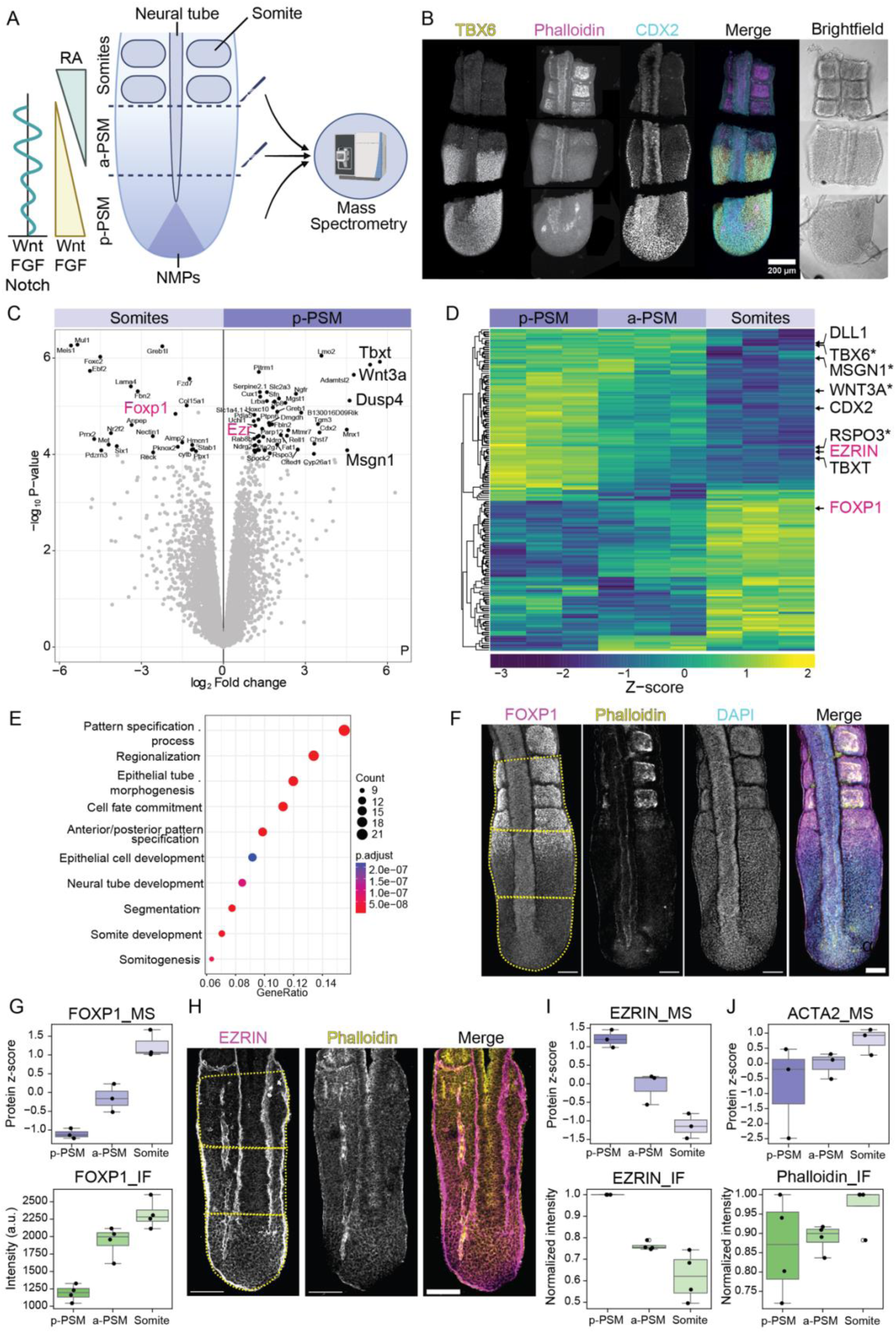
Spatial proteomics of mouse embryo tails reveals tissue-wide protein expression patterns. **A** Schematic representation of embryonic tail with oscillations and morphogen gradients regulating somitogenesis (*left)* and experimental procedure (*right)*: Embryonic tails were dissected into posterior tissue including posterior PSM (*p-PSM*), anterior tissue including anterior PSM (*a-PSM*) and tissue including two last 2 formed somites (*Somites*). Eight tails pooled per sample and 3 independent experiments. **B** Representative image of dissected embryonic tail stained with antibodies against TBX6 (maximum intensity projections, MIP), CDX2 (single-z plane) and counterstained with Phalloidin (MIP) and brightfield (single-z plane). Dissections performed in a stereoscope based on the brightfield. Scale bar 200 µm. **C** Volcano plot showing proteins differentially expressed between p-PSM and Somites regions. Selected new hits are highlighted (magenta). **D** Clustermap displaying differentially expressed proteins between any of the regions. Selected known proteins (black) and new hits of interest (magenta) are annotated. * indicates proteins with 1-3 imputed values. Complete annotated clustermap in Supplementary Data 1. **E** GO term enrichment analysis of the 148 differentially expressed proteins found in proteomics dataset. **F, G**: Validation of FOXP1 protein expression in embryonic tails. **F** Representative images of FOXP1 immunostainings in mouse embryonic tail counterstained with DAPI and Phalloidin. Scale bar 100 µm. **G** Mass-spectrometry results of FOXP1 (FOXP1_MS) and quantification of FOXP1 intensity levels measured in immunostainings in the regions of interest highlighted with yellow dashed lines in **F** (FOXP1_IF, n = 4). **H-J**: Validation of Ezrin expression pattern in embryonic tails. **H** Representative images of EZRIN immunostainings in embryonic tails counterstained with Phalloidin. Scale bar 100 µm. **I** Mass-spectrometry results for EZRIN (EZRIN_MS) compared to quantification of Ezrin in immunostainings in the regions of interest highlighted with yellow dashed lines in (H), (EZRIN_IF, n = 4). **J** Mass-spectrometry results of Actin alpha 2 levels (ACTA2_MS) compared to quantification of Phalloidin (filamentous Actin) stainings in regions of interest highlighted in yellow dashed lines in H (Phalloidin_IF, n = 4).

Despite advances in our understanding of somitogenesis, several questions remain unresolved regarding the detailed mechanisms modulating signalling dynamics along the AP axis and somite formation. To address these questions, the cellular components along the PSM must be identified. Transcriptomic studies have allowed the identification of cyclic or graded expression of many genes in somitogenesis^26–30^. However, previous studies aiming at identifying cyclic genes focussed on posterior PSM^20,27^, while somite formation occurs in the anterior PSM. In addition, the majority of studies so far focussed on mRNA levels in the embryo rather than analysing protein levels^20,26–28^. However, in most cases, it is the protein product that carries out the gene’s function and protein and mRNA levels often do not correlate (e.g.^31,32^). While protein dynamics have been shown for selected cyclic genes in somitogenesis, such as

Dll1, Notch1, Hes1 and Hes7^33–36^, levels and dynamics of many other proteins remain unknown. Here, we present a systematic, multi-omics approach to dissect the proteome of mouse somitogenesis with spatial and temporal resolution. We first used mass-spectrometry on dissected mouse embryonic tail tissue to map spatial protein expression and compared it to mRNA levels obtained by RNA sequencing. To capture dynamic changes, we developed a microfluidic platform that synchronizes signalling oscillations across multiple non-adherent tail explants. Combining this synchronization system with transcriptomic and proteomic analysis enabled the identification of oscillatory mRNAs and proteins in mouse somitogenesis. Leveraging this dataset, we uncovered a novel mechanism in which dynamic, antagonistic gradients of ligands and receptors regulate Wnt signalling along the embryonic anteroposterior axis.

## Results

### Spatial proteomics identifies differential protein patterns in segmenting mouse embryo tails

Before investigating oscillatory proteins in somitogenesis, we first mapped the spatial proteome of mouse embryo tails along the AP axis. To this end, mouse tails from embryonic day 10.5 (E10.5) were manually dissected based on morphology: posterior (includes posterior PSM, NMPs, neural tube and skin, Tbx6/Cdx2-high region), anterior (includes anterior PSM, neural tube, gut and skin, Tbx6 high/ Cdx2 low-region) and somites (two most-recently formed somites, neural tube, gut, skin, and other cell types) hereafter called p-PSM, a-PSM and somites regions, respectively (Fig. 1A, B). To minimize protein degradation, dissection was limited to whole regions without further dissection. Eight samples were pooled to improve low-abundance protein detection and average segmentation clock phase differences between embryos.

Label-free quantification (LFQ) mass spectrometry identified between 7000 and 8000 unique proteins (Extended Data Fig.1A, Supplementary Data 1). To assess sample reproducibility and variability, we used principal component analysis (PCA). PC1 separated samples based on their location along the AP axis from somites to a-PSM and p-PSM regions (Extended Data Fig.1B). Despite the presence of other tissues, most of the dissected tissue corresponds to PSM and somites (Fig. 1B). Correspondingly, we can detect PSM-specific proteins in our dataset such as TBX6 (Extended Data Fig.1C, D), confirming that the dataset captures the proteome of mouse somitogenesis.

We next searched for spatially distinct protein expression through pairwise tissue region comparisons. Overall, when compared to somites, p-PSM and a-PSM showed an upregulation of known PSM-related proteins (Fig. 1C, Extended Data Fig. 1C). Between p-PSM, a-PSM and somites, 148 proteins were significantly differentially expressed (Fig. 1D), accounting for 2% of the total dataset. Conversely, most detected proteins showed no significant differential expression, highlighting the specificity of our approach in capturing spatial expression patterns. Among the differentially expressed proteins were the known PSM markers WNT3A, MSGN1, TBXT CDX2, TBX6, and DLL1 (Fig. 1D, Supplementary Figure 1), which displayed the expected expression patterns^14,37–39^ (Extended Data Fig.1E-G). In contrast, among proteins not displaying a differential expression pattern was TUBA4A, encoding for alpha-tubulin, a core component of microtubules (Extended Data Fig.1H-J). Thus, we could identify known graded proteins related to somitogenesis.

We then focussed on new differentially expressed proteins that might be related to somitogenesis. GO term analysis on differentially expressed proteins revealed that processes including segmentation, differentiation and AP pattern specification were enriched (Fig. 1E). One differentially expressed protein was FOXP1, a transcription factor broadly expressed in zebrafish embryos^40^ and implicated in neurodevelopmental disorders in humans^41^. We found high FOXP1 expression in somites, intermediate expression in a-PSM and low in p-PSM (Fig. 1D), which we confirmed by immunostaining of E10.5 embryonic tails (Fig. 1F, G). Moreover, EZRIN, part of the Ezrin-Radixin-Moesin (ERM) proteins linking the plasma membrane to underlying actin cytoskeleton^42^, displayed a posterior-to-anterior gradient (Fig. 1D), which we validated by immunostaining and quantification of EZRIN and counterstaining for F-Actin in tissue sections of E10.5 mouse tails (Fig. 1H-J). This shows that our dataset can identify differentially expressed proteins in the PSM, even if they are also present in other tissues. By mapping the spatial proteome of the segmenting mouse embryo tail, we uncover PSM-associated proteins and graded proteins that may be relevant for somite formation.

### mRNA and protein expression patterns do not always correlate in developing embryos

Most insights into embryonic development stem from mRNA analyses, as ISH probes are generally easier to generate than specific antibodies, and RNA-sequencing is more sensitive and requires less material than mass spectrometry. To determine whether protein expression correlates with transcript expression during somitogenesis, we performed bulk RNA-sequencing on p-PSM, a-PSM and somites regions (akin to Fig. 1A). PCA analyses showed tight clustering of triplicates from the same regions (Extended Data Fig. 2A). We identified ∼18,000 unique transcripts with 2,589 (14%) differentially expressed genes (Extended Data Fig. 2B, Supplementary Data 2). Among these were somitogenesis-related genes (e.g., *Mesp2*, *Uncx*, *Tbxt*) and to a smaller extent from other lineages, such as *Neurog1* for neurogenesis.

To assess transcript–protein expression correlation, we calculated Spearman’s Rho (r) correlation scores between z-scores of each protein–mRNA pair in both the total protein and differentially expressed protein datasets (Fig. 2A, Extended Data Fig. 2C). By comparing z-scores, we compared protein and mRNA expression patterns to each other rather than absolute levels. Many protein patterns correlated well with the corresponding transcripts, including PSM- and somitogenesis-related genes such as *Wnt3A, Dusp4* and *Dll1* (r > 0.5: 3066 out of 7007 protein-mRNA pairs (43.8 %), Fig.2A, B). In contrast, others showed a low correlation such as *Itga1* (−0.5 > r < 0.5: 2041 protein-mRNA pairs (29.1 %), Fig. 2A, B) or even anti-correlation such as *Pxdn* (r < −0.5: 1900 protein-mRNA pairs (27.1 %), Fig. 2A, B). As a validation, we performed a staining for *Pax3* mRNA and protein, whose expression patterns with increasing levels from p-PSM towards the somite region were predicted to be highly correlated (r = 0.78) (Extended Data Fig.2D).

**Figure 2.**
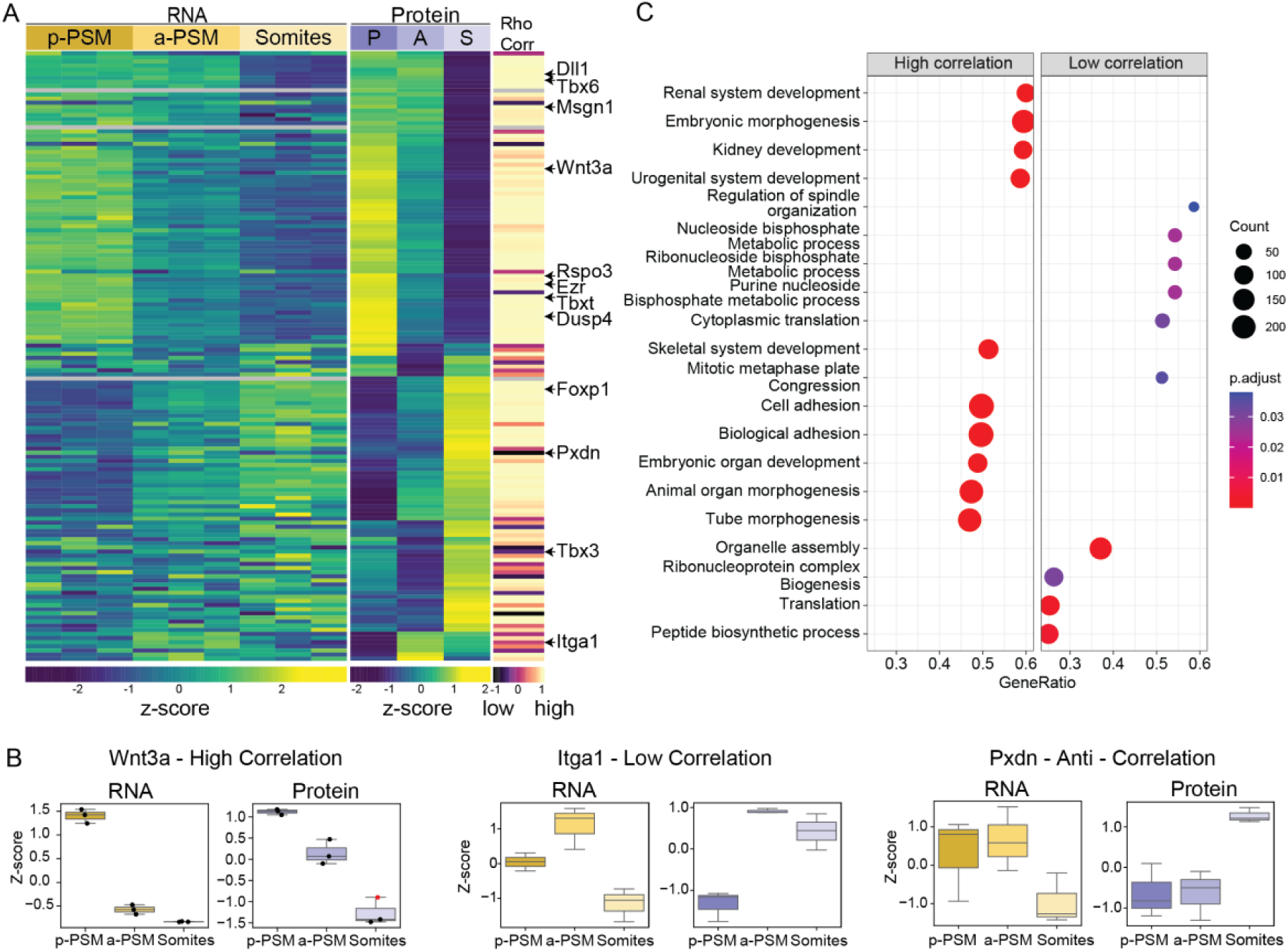
Correlation between Protein and RNA expression. **A** *Left panel*: Heatmap of transcripts (data from performed RNA-seq) corresponding to the differentially expressed proteins. *Middle panel*: Heatmap of differentially expressed proteins (corresponds to Fig. 1D averaged per region, p-PSM (P), a-PSM (A), Somites (S)). *Right panel*: colour gradient to visualize Spearman’s Rho correlation score for each protein-RNA pair. Selected genes of interest are annotated. **B** Selected examples (from A) of transcripts and proteins with high, low, or anti – correlation Spearman’s score. For Wnt3a, a value was imputed, single values are indicated and imputed value is highlighted in red. **C** Top 20 enriched gene sets identified by gene set enrichment analysis (GSEA) of genes ranked by Spearman correlation between RNA and protein expression levels across Somites, p-PSM and a-PSM.

To link RNA–protein correlation patterns to cellular pathways, we performed GSEA analysis on the total dataset (7,007 protein-mRNA pairs, Fig. 2C). Highly correlated pairs were liked to embryonic morphogenesis, cell adhesion and organ morphogenesis, while low correlation to metabolic processes, translation and ribonucleoprotein biogenesis (e.g. Wnt, Notch, FGF signalling and metabolic enzymes in Extended Data Fig. 2E-I).

Thus, while many genes show high RNA–protein correlation, others diverge, underscoring the importance of protein-level analysis for functional investigation of somitogenesis.

### Establishing a microfluidic system to synchronize genetic oscillations in embryonic tails

Since somitogenesis is a dynamic process, we next applied time-resolved omics to identify oscillating transcripts and proteins. As pooling tails from multiple embryos was needed for low abundance protein detection, synchronizing samples to the same segmentation clock phase was crucial. To this end, we established a microfluidic system based on our previous work^21,32^ to enable the culture of non-adherent growing embryonic tail explants in 3D.

E10.5 mouse embryo tails expressing the *lunatic fringe* (*Lfng*) reporter LuVeLu^17^ were cultured on-chip (Fig. 3A-C, Supplementary Video 1). Pulses of either DMSO (control) or the Notch signalling inhibitor DAPT (N-[N-(3,5-difluorophenacetyl)-l-alanyl]-S-phenyl glycine t-butyl ester), a gamma-secretase inhibitor, were applied with a period of 130 min (Fig. 3A-C, Extended Data Fig. 3A-B). Entrainment with DAPT led to synchronization of LuVeLu oscillations in anterior and posterior PSM after approximately 5 pulses, while entrainment with DMSO did not (Fig. 3D, Extended Data Fig. 3B). We confirmed that Wnt signalling was successfully entrained as well, using tails expressing the Wnt signalling reporter Axin2-T2A-Venus (Axin2T2A)^18^ (Extended Data Fig. 3C-E). Interestingly, in entrained tails, Notch- and Wnt-signalling waves showed defects: After 3 – 4 drug pulses, anterior PSM cells skipped an oscillation compared to posterior cells (Fig. 3E, Extended Data Fig. 3C). After up to 6 oscillation cycles this was resolved with waves travelling along the entire length of the PSM.

**Figure 3.**
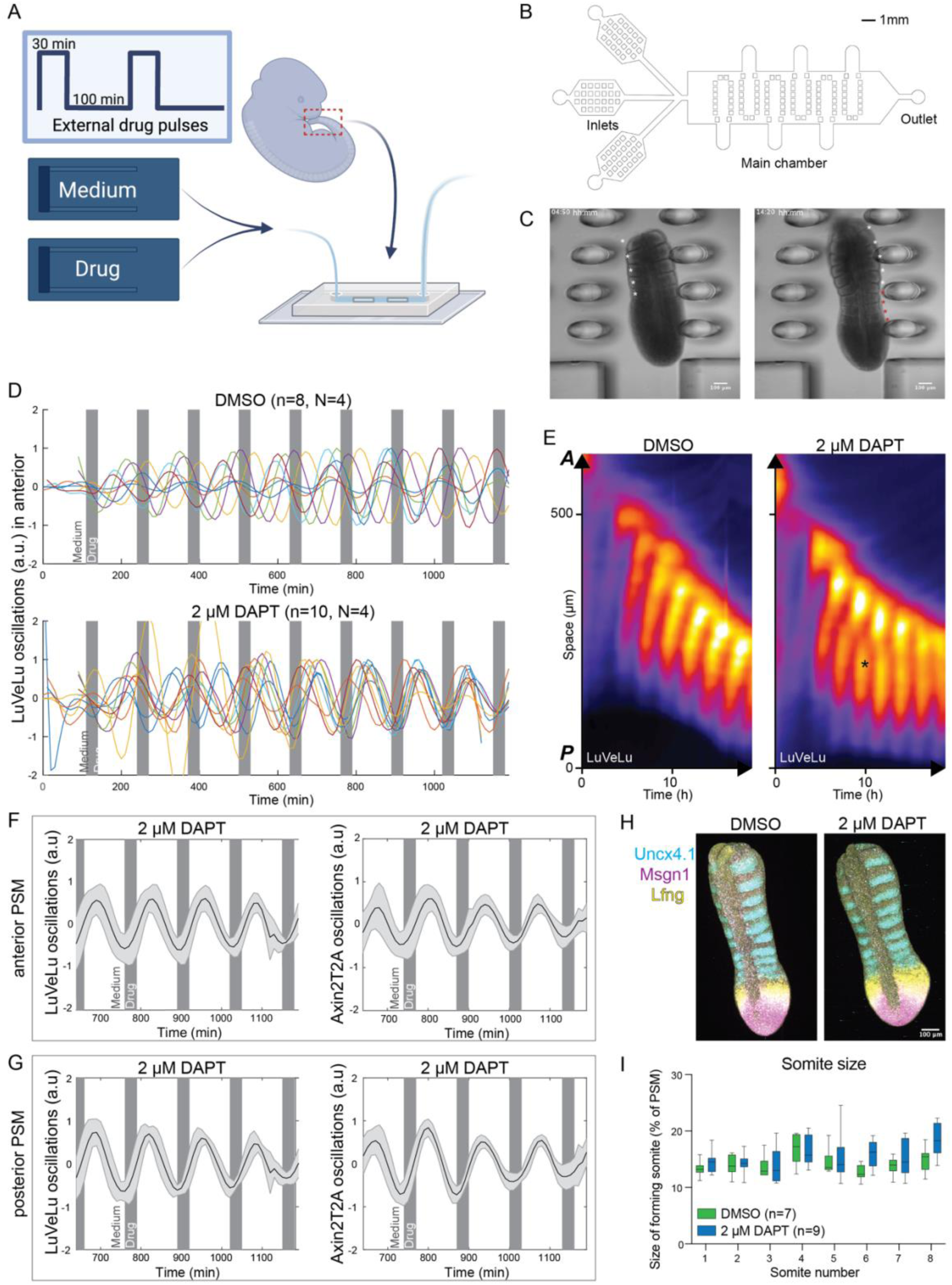
Microfluidic entrainment of 3D growing mouse embryo tails. **A** Experimental overview: E10.5 mouse embryo tails were dissected, loaded into a microfluidic chip and entrained with 100 min medium pulses and 30 min pulses of medium + drug (DMSO or DAPT). **B** Microfluidic chip design. Tails are entrapped with pillars within the main chamber. **C** Representative brightfield images of tail growing during a microfluidics experiment. *Left panel*: t1 = 04:50 hh:mm. Right panel: t2 = 14:30 hh:mm. White asterisks indicate somite boundaries present at t1, red asterisks indicate newly formed boundaries at t2. **D** Detrended LuVeLu signal in anterior PSM (quantified from kymographs as described in Extended Data Fig.3A). Gray vertical bars in plots indicate moment of medium / drug pulsing. **E** Representative kymographs of LuVeLu oscillations after entrainment with DMSO (*left*) or DAPT (*right*). Skipped oscillation in DAPT treatment highlighted with asterisk. **F, G** Average and standard deviation of oscillations of Notch signalling using LuVeLu reporter (*left panel*) and Wnt signalling using Axin2T2A reporter (*right panel*) in anterior **(F)** and posterior PSM **(G)** upon entrainment with DAPT. **H** Representative images of control and entrained tail explants stained with markers for rostral somites (*Uncx4.1*), posterior PSM (*Msgn1*) and the segmentation clock (*Lfng*) using HCR. **I** Quantification of somite size in control (DMSO) and entrained (DAPT) LuVeLu-expressing tail explants properly positioned in real-time imaging experiments (panel D) (unpaired T test not significant, N=3). Scale bars 100 µm.

Upon entrainment, the in-phase relationship between Wnt- and Notch-signalling oscillations in anterior PSM was maintained (Fig. 3F), as previously shown to be necessary for proper somite formation^18^. Unlike previous entrainment experiments using 2D adherent PSM cultures^18^, Wnt- and Notch-signalling oscillations in posterior PSM also approached an in-phase relationship (Fig. 3G). In agreement, the velocity of both LuVeLu and Axin2T2A waves increased (Extended Data Fig.3F, G). Nevertheless, tissue growth, somitogenesis and differentiation continued throughout the course of the experiment (Fig. 3H-I).

Thus, we have established a microfluidic system that enables the synchronization of signalling oscillations in embryonic tails, while maintaining tissue growth and somite formation in a 20-hr experiment.

### Spatiotemporal proteomics to investigate segmentation clock dynamics

To capture dynamic expression at both the protein and RNA levels, we combined our microfluidic entrainment method with omics analysis. To allow for fast tissue handling, we adapted the inner chamber of the chip to be able to simultaneously load and recover multiple tails (Fig. 4A). The overall dimensions of the chip were maintained to ensure equivalent liquid flow and drug exchange between the two chip designs. Wild-type embryonic tails (E10.5) were entrained with a period of 130 minutes. Experiments were terminated with 30-minute intervals throughout one oscillation cycle (30, 60, 90, and 120 minutes following the sixth drug pulse, Fig. 4B), after which embryonic tails were dissected into p-PSM, a-PSM and somite regions (Fig. 4C). Samples from four tails per region and timepoint were pooled, with quadruplicates for proteomics and triplicates for RNA-sequencing. PCA showed both protein and RNA separated by tissue region in PC1 (Extended Data Fig.4A, B). After filtering for proteins/mRNAs with at least two replicates for all timepoints for each region and not having more than 6 values missing in total, ∼7,600 proteins and ∼14,000 transcripts remained (Extended Data Fig.4C).

**Figure 4.**
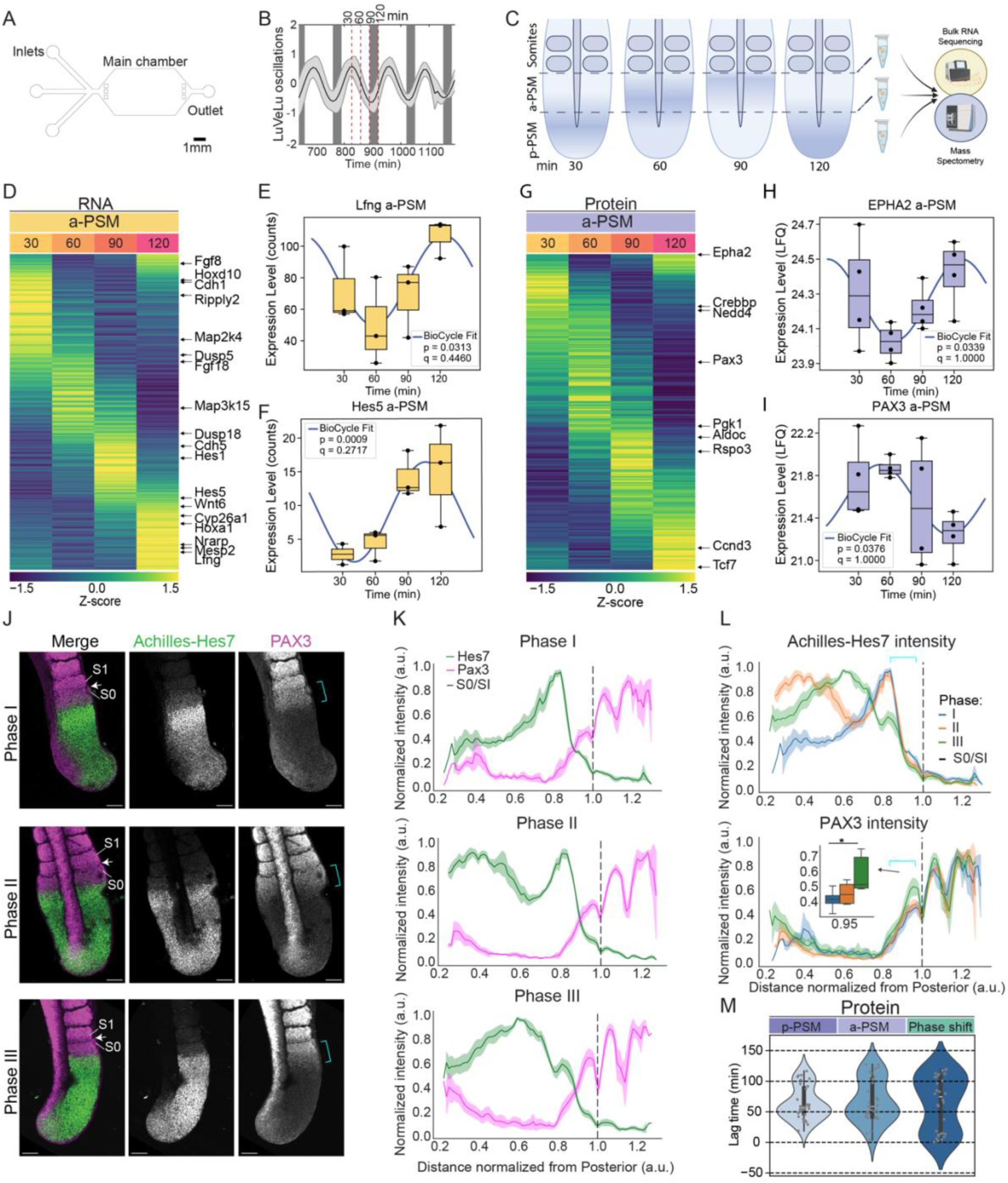
Spatiotemporal proteomics and bulk RNA-seq reveal protein/transcripts dynamic expression in somitogenesis. **A** Microfluidic chip design for experiments combining entrainment with subsequent omics analysis. **B, C** Experimental design: **B** E10.5 wild-type tail explants were entrained with a 130 min period (100 min medium + 30 min drug). Average oscillation dynamics of Notch signalling using LuVeLu reporter (same as in Fig. 3F) overlaid with dashed red lines illustrating the 4 timepoints used for omics analysis. **C** Embryonic tails were dissected into p-PSM, a**-**PSM and somite regions and samples from 4 embryos were pooled. **D** Heatmap of significantly dynamic transcripts (p<0.05) in a-PSM. Genes ordered by lag time (time between oscillating gene peaking and a sine wave peaking at 0 min). Selected known cyclic transcripts and new dynamic ones are annotated. **E, F** Boxplots of mass-spectrometry data of examples of significantly dynamic transcripts (p<0.05) detected in a-PSM, segmentation clock gene *Lfng* (**E**) and *Hes5* (**F**). Fitted sinusoidal from BYO_CYCLE analysis is overlaid and p and q values are indicated. **G** Heatmap of significantly dynamic proteins (p<0.05) in a-PSM. Proteins ordered by lag time. Selected proteins are annotated. **H, I** Boxplots of mass-spectrometry data of examples of significantly dynamic proteins (p<0.05) detected in a-PSM, EPHA2 (**H**) and PAX3 (**I**). Fitted sinusoidal from BYO_CYCLE analysis is overlaid and p and q values are indicated. **J** Immunostaining of PAX3 in E10.5 mouse embryonic tails expressing Achilles-Hes7, separated into 3 distinct phases. Arrows point to somite boundary between the newly forming somite (S0) and the previously formed somite (S1). Cyan brackets: PAX3 expression upregulation in a-PSM and S0. Scale bar 100 µm. **K** Normalized intensity of Achilles-Hes7 and PAX3 quantified from posterior until S3 in each phase. Dashed line x=1.0 marks S0/S1 boundary (n=12, N=5), same as arrows shown in **J**. **L** Normalized Achilles-Hes7 (*top*) and PAX3 (*bottom*) intensity per phase. Cyan brackets: location of S0 as shown in **J.** PAX3 expression in a-PSM at **x=**0.95 highlighted in inset boxplot (* p<0.05, unpaired T-Test, two-sided). **M** Violin plot of lag time for significantly dynamic proteins (p<0.05) in p-PSM and a-PSM and phase-shift between the two regions.

To identify dynamic expression, we used BIO_CYCLE^43^, a neural network-based analysis which identifies oscillatory expression in omics data by fitting periodic models using a Bayesian framework. This captures rhythmic patterns, such as circadian or ultradian rhythms, even in the presence of biological noise or missing data. We trained separate neural networks for protein and RNA using posterior datasets and analysed the different regions to predict oscillating proteins and transcripts (using Hes7, Lfng, Dusp4, Dusp6, Axin1 and Axin2 as ground truth, if present in the dataset). Transcripts and proteins rendered significantly cycling based on BIO_CYCLE, could either reflect oscillations within single cells, as observed for e.g., *Hes1* or *Hes7*^34,35^, or dynamic expression patterns on tissue level, as observed for expression of the differentiation factor *Mesp2* in anterior PSM, which occurs once in a given cell^3^. Only repeated measurements in the same cell by for example real-time imaging of reporter constructs could resolve this. We will therefore refer to all hits as “significantly dynamic” rather than “significantly oscillating”.

We identified 495, 311, and 253 dynamic proteins (p < 0.05) in p-PSM, a-PSM, and somites, respectively, and 989, 1084, and 1339 dynamic mRNAs (Extended Data Fig. 4C–E). BIO_CYCLE estimates oscillation-peak times relative to a reference sine wave peaking at 0 min, termed lag time^47,48^. We visualized significantly dynamic data in heatmaps ordered by lag time (Fig. 4D-I, Extended Data Fig. 4F-I). While most dynamic hits did not pass multiple-testing correction (q > 0.05), several known segmentation clock genes showed nominal significance (p < 0.05), including *Hes7* and *Hey1* (mRNA p-PSM, Extended Data Fig.4F), *Lfng* and *Hes5* (mRNA a-PSM, Fig. 4E, F), DUSP6 (protein p-PSM, Extended Data Fig.4G) and EPHA2 (protein a-PSM, Fig. 4H). These results support our combined microfluidics-omics approach in identifying candidate dynamic genes and proteins.

GO term analysis indicated that in p-PSM, many detected dynamic proteins were related to Wnt signalling and patterning but also metabolic and catabolic activities (Extended Data Fig.4J, K). In a-PSM, dynamic transcripts and proteins were associated with neural tube development, ribosomal processes and epigenetic regulation (Extended Data Fig.4J, K). In somites, many dynamic transcripts and proteins belonged to morphogenetic processes such as epithelial tube and limb morphogenesis but also to stress response (Extended Data Fig.4J, K). Some notable proteins predicted to be dynamic were components of the electron transport chain such as cytochrome c oxidase 2 COX2 and ATP synthase subunit ATP5F1A, the glycolytic enzyme ALDOC or the centrosomal protein Cep152 (Extended Data Fig.4F-I, Supplementary Information).

*Pax3*, a transcription factor associated with dermomyotome differentiation and expressed in somites and neural tube^44^ (MAMEP database http://mamep.molgen.mpg.de/), was predicted as dynamic protein in a-PSM (Fig. 4I). To validate its dynamics, we performed immunostaining for PAX3 in Achilles-Hes7-expressing E10.5 mouse embryo tails (Fig. 4J). Based on the Achilles-Hes7 expression pattern, we classified embryonic tails into three distinct phases of the segmentation clock (Fig. 4J, K, Extended Data Fig. 5A-D, Supplementary Video 2). Phase I shows peak expression in anterior PSM, just posterior to the newly forming somite (S0); in Phase II, anterior expression declines while a new wave emerges in posterior PSM; Phase III is marked by broad intermediate expression across the PSM and formation of a new somite. PAX3 was expressed in anterior PSM around S0 and in somites in all phases (Fig. 4K, L). When grouping the data by Achilles-Hes7 phases, we found that PAX3 expression was broader in anterior PSM during Phases I and II but became narrower with a distinct stripe at S0 in Phase III (cyan bracket in Fig. 4L), indicating dynamic protein expression likely linked to periodic somite differentiation.

Plotting the expression profiles (z-score) of detected dynamic transcripts and proteins ordered by their lag time revealed gene clusters peaking at distinct timepoints within the oscillation cycle (Fig. 4D, G, Extended Data Fig. 5E, F, 4F-I). The average period provided by BIO_CYCLE was approximately 129.9 min (protein a-PSM), consistent with the mouse segmentation-clock period^45^. Proteins mainly peaked at ∼50 min and to a lesser extent at ∼110 min in p-PSM and a-PSM (Fig. 4M), indicating that many proteins peak out-of-phase relative to each other. Transcripts had a similar distribution of lag times with a main peak at ∼110 minutes and a second one at ∼30 min (Extended Data Fig.5G). Signalling waves travel with different speeds through the PSM^14,18^, which should result in a phase-shift (hence different lag times) between posterior and anterior. We calculated the phase-shift between p-PSM and a-PSM for transcripts or proteins detected as significantly dynamic in both regions (see Methods for details, Extended Data Fig. 5E,F). The phase-shift distributions for both transcripts and proteins showed two peaks: at ∼20 and ∼100 min (Fig. 4M, Extended Data Fig. 5G), indicating different wave velocities for different transcripts and proteins. This dataset can therefore provide interesting indications on wave dynamics and phase-relationships of dynamic genes for future investigations.

Our microfluidics-integrated omics approach captured dynamic expression of known clock components and identified candidate oscillatory genes and proteins.

### Dynamic antagonistic receptor-ligand expression modulates oscillation parameters

Morphogen gradients regulate somitogenesis, including that of *Wnt3a* with high expression in posterior and decreasing towards the anterior PSM^14,20,38^, a spatial gradient we also found at protein level in our mass-spectrometry data (Fig. 1, 2). Additionally, Wnt signalling is part of the segmentation clock^14,17,18^. While the ligand peaks in posterior PSM/ tailbud, Wnt-signalling oscillations increase in amplitude and intensity towards the anterior PSM^14,18^ (Fig. 5A). How this apparent discrepancy can be reconciled remains unknown. To address this, we analysed expression of Wnt signalling components in our datasets. Interestingly, while ligands were highest in p-PSM, receptors were mostly detected in a-PSM or somite regions (Fig. 5B). It is noteworthy that we observed this antagonistic ligand-receptor expression not only for Wnt signalling, e.g., *Wnt3A* and *Fzd1/2/7*, but also for other pathways such as FGF and Notch signalling (Extended Data Fig.6A, B).

**Figure 5.**
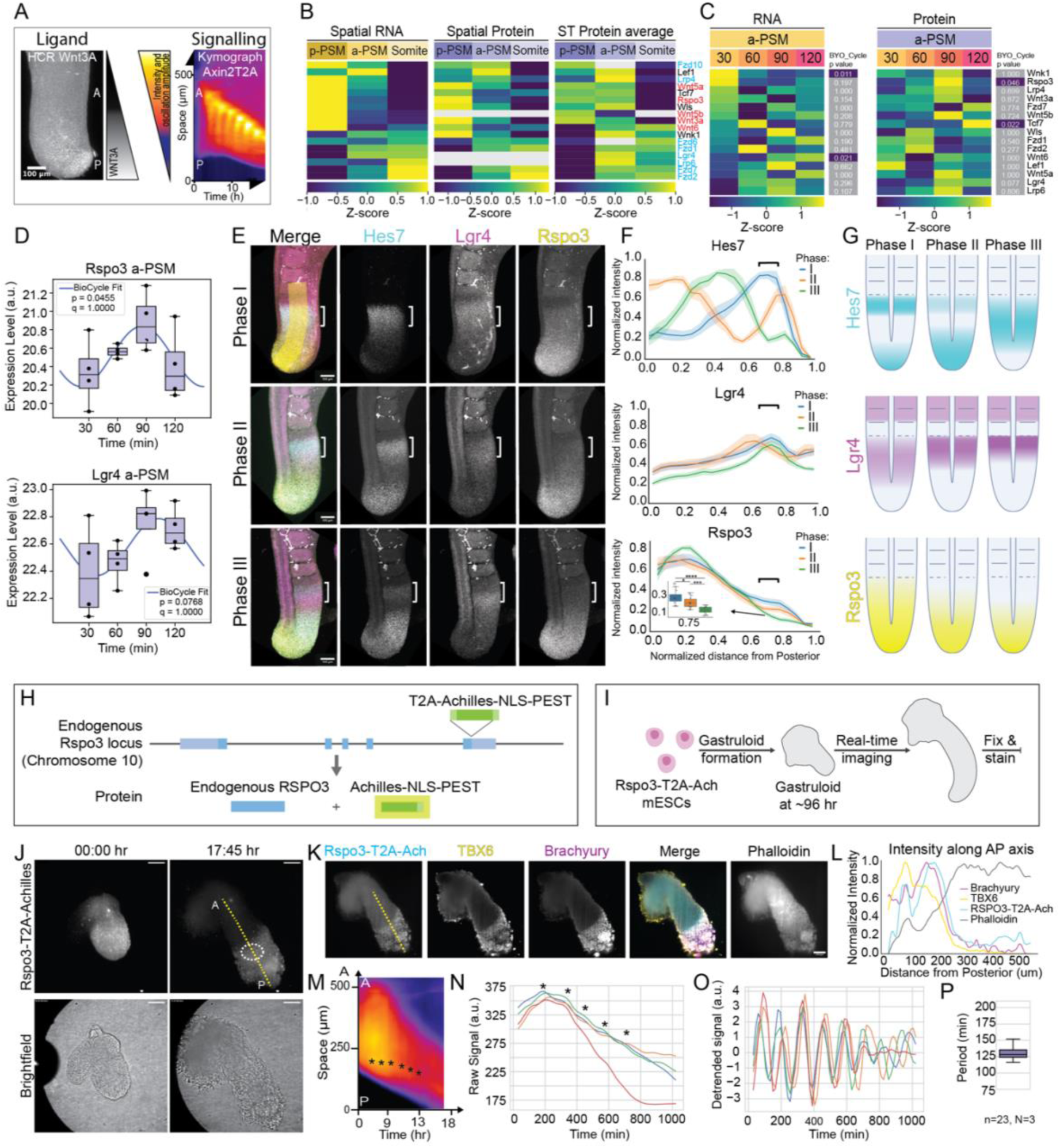
Dynamic antagonistic receptor-ligand expression modulates signalling dynamics. **A** Ligand *Wnt3a* HCR in E10.5 mouse embryonic tail (*left*). Kymograph of embryonic tail expressing AxinT2A Wnt signalling reporter (*right*). **B** Heatmap of selected Wnt-signalling components in spatial RNA-seq (*left*), spatial proteomics (*middle*) and averaged timepoints for each region of spatiotemporal (ST) data (*right*). Average per region is shown for all datasets. (Ligands: red, receptors: cyan). **C** Heatmap of selected Wnt-signalling components found in spatiotemporal RNA and protein data in a-PSM and corresponding p-values from BIO_CYCLE analysis. Highlighted in blue: significant values (p<0.05). **D** Boxplots of mass-spectrometry data for RSPO3 (ligand) and LGR4 (RSPO3 receptor) in a-PSM. Fitted sinusoidal from BYO_CYCLE analysis is overlaid and p and q values indicated. **E** Representative images of staining for *Hes7*, *Lgr4* and *Rspo3* mRNA using HCR in E10.5 mouse embryo tails, categorized in 3 distinct phases according to *Hes7* oscillations. Brackets point to future somite. **F** Quantification of *Hes7*, *Lgr4* and *Rspo3* intensity for each phase. Intensity quantified along a line (shown in yellow in **E)** from posterior tip to S0/S1 boundary and normalized to the maximum intensity per tail (mean + SEM). Brackets: future forming somite as in **E**. *Rspo3* expression in anterior PSM at x=0.75 in inset boxplots. **G** Schematic summarizing *Hes7*, *Lgr4* and *Rspo3* dynamic expression relative to the Hes7 oscillation phase. **H** Schematic of genome editing of the *Rspo3* locus to insert a cassette expressing T2A-Achilles-NLS-PEST. **I** Experimental procedure of gastruloid formation, real-time imaging and immunostaining. **J** Representative images of a gastruloid at the beginning (t=0h) and end (t=17h45) of real-time live-cell imaging. P: Posterior, A: Anterior pole. **K, L** Fixed gastruloid after live cell imaging (same as in **J**) immunostained for TBX6 and T/Brachyury and counterstained with Phalloidin (**K**). **L** Quantification of intensity along a line from posterior to anterior regions in the gastruloid (yellow dashed line in **J**). **M** Kymograph of Rspo3-T2A-Achilles intensity along a line from posterior to anterior (corresponding to yellow dotted line in **J**). Asterisks: vertical lines in the kymograph indicating oscillations. **N, O** Quantified raw (**N**) and detrended (**O**) signal from four example gastruloids from a single experiment, using a static ROI in the positive Rspo3-T2A-Achilles region (dotted oval ROI in **J**, corresponding to anterior PSM). Asterisks in **N** highlight oscillation peaks. **P** Quantification of Rspo3-T2A-Achilles period (n=23, N=3). Scale bars 100 µm.

To understand the significance of this antagonistic expression, we focussed on R-Spondin/LGR signalling, known to amplify Wnt signalling^46–52^. When R-SPONDIN binds to its receptor LGR (leucine-rich repeat-containing G-protein-coupled receptor), this complex sequesters the E3 ligases RNF43 or ZNRF3 leading to the stabilisation of Wnt receptors and amplification of Wnt signalling^46,49^. In our mass-spectrometry data, R-SPONDIN 3 (RSPO3) was highest in p-PSM (Fig. 5B), as previously shown at mRNA level^26,28,53^. In contrast, its receptor LGR4 was highest in a-PSM (Fig. 5B). To validate the expression patterns of Rspo3 and Lgr4, we performed ISH in E10.5 tails. *Rspo3* showed a posterior-to-anterior gradient, while *Lgr4* was enriched in the anterior PSM (Extended Data Fig.6C, D), confirming the antagonistic ligand–receptor pattern. ISH of *Rnf43*^54^ indicated a similar expression band in anterior PSM as *Lgr4* (Extended Data Fig.6C). In contrast, other LGR or RSPO isoforms showed no specific expression patterns (data not shown).

Notably, several Wnt signalling components including RSPO3 were deemed dynamic in our dataset (Fig. 4D, G, Extended Data Fig. 4F-I, 5C, D; RSPO3: p = 0.046). To confirm *Rspo3* dynamics in anterior PSM, we used hybridization chain reaction (HCR) to correlate mRNA expression of *Rspo3* and *Lgr4* to that of the known cyclic gene *Hes7*. We used the established oscillation-phase classification for Achilles-Hes7 to separate tails based on *Hes7* HCR into three oscillation phases (Fig. 4J-L, Extended Data Fig. 5). *Rspo3* showed a posterior-to-anterior gradient in all phases, but its expression in anterior PSM changed with Hes7-oscillation phases (Fig. 5D-F, brackets): In Phase I and II, the *Rspo3* gradient reached the future somite boundary, where a sudden drop in signal was visible (Fig. 5D-F); in Phase III, the gradient was more restricted to the posterior PSM (Fig. 5E-G). In addition, in anterior PSM expression of *Rspo3* overlapped with the *Hes7* stripe in Phase I and decreased in intensity in the following Phases II and III as Hes7 expression also declined (Fig. 5E-F brackets). *Lgr4* also showed a dynamic expression pattern during the *Hes7* oscillation cycle (Fig. 5E-G, Extended Data Fig. 6E). The anterior stripe of *Lgr4* expression did not overlap with the bright *Axin2* stripe in forming somites but with the dynamic *Axin2* expression in anterior PSM. These analyses indicate a dynamic expression of both *Rspo3* and *Lgr4* in anterior PSM during somitogenesis.

Since RSPO3 was predicted to be dynamic at the protein level, we generated a mouse embryonic stem cell (mESC) reporter line. To this end, we endogenously tagged *Rspo3* with the fluorescent protein Achilles, a nuclear localization signal (NLS) and PEST degradation sequence, separated from the *Rspo3* gene by a T2A sequence, a ribosome-skipping site leading to the formation of two independent proteins^34,55,56^ (Fig. 5H, Extended Data Fig.6F). This approach enabled visualization of RSPO3 expression and translation dynamics without disrupting protein function. We used reporter mESCs to generate gastruloids, *in vitro* embryo-like structures recapitulating the three germ layers in post-gastrulation embryos, axis elongation and segmentation-clock oscillations^57^. At 96 h, gastruloids were embedded in 10% Matrigel, imaged for ∼18 h in a light-sheet microscopy, fixed and stained for PSM markers TBX6 and T (Fig. 5I-L). Gastruloids displayed graded RSPO3 reporter expression with high levels in the posterior pole (defined by T and TBX6 staining, Fig. 5J-L). Real-time imaging revealed that the gradient regressed in a stepwise manner towards the posterior, as segments started to appear in the anterior region (Fig. 5J, M, Supplementary Video 3). Analysing the Rspo3-T2A-Achilles expression in a region corresponding to anterior PSM over time, we indeed detected oscillations with low amplitude (Fig. 5N, O, Extended Data Fig. 6G). RSPO3-reporter oscillations had a period of ∼130 min (Fig. 5P, Extended Data Fig. 6H-J), corresponding to the mouse segmentation clock period^45^. Thus, RSPO3 is expressed in a gradient along the AP axis of the PSM with oscillatory expression in anterior PSM, where its receptor *Lgr4* is also dynamically expressed.

Next, to reveal the function of the dynamic expression pattern of RSPO3-LGR4, we perturbed the RSPO3 by either adding exogenous RSPO3 in excess or by preventing the interaction of RSPO3 with its receptor by adding the extracellular domain of LGR5 (LGR5-ECD) as soluble protein previously shown to sequester extracellular RSPO^46,49^ (Fig. 6A, B). Addition of RSPO3 and LGR5-ECD had the expected effect on Wnt-signalling activity in HEK293 Super TOPFlash (STF) reporter cells^49^ (Extended Data Fig.7A). We then perturbed RSPO-LGR in mouse PSM 2D spread-out cultures expressing the Wnt-signalling reporter Axin2T2A^18^ and performed real-time imaging (Supplementary Video 4). In these 2D cultures, the AP axis re-organizes with signalling waves travelling in a centre-to-periphery direction and cells spreading towards the periphery^45^ (Fig. 6B), which we reasoned would ensure high penetrance of external proteins into the tissue. Surprisingly, both RSPO3 and LGR5-ECD treatments reduced Axin2T2A intensity and oscillation amplitude in anterior PSM without altering the period (Fig. 6C-F). Axin2 is expressed not only in the PSM but also as a stable stripe in newly formed somites, reflecting differentiation and AP patterning (Fig. 6B). Consistent with previous studies^14,58^, this Axin2 somite stripe seemed to be regulated by Notch rather than Wnt signalling (Extended Data Fig.7B). Interfering with RSPO3 dynamics markedly reduced Axin2T2A stripe intensity in forming segments, exceeding the decrease observed in the PSM (Fig. 6G, H). This reduction appeared to reflect not only lower Axin2 intensity per cell, but also fewer cells with elevated Axin2 levels, together suggesting impaired somite differentiation.

**Figure 6.**
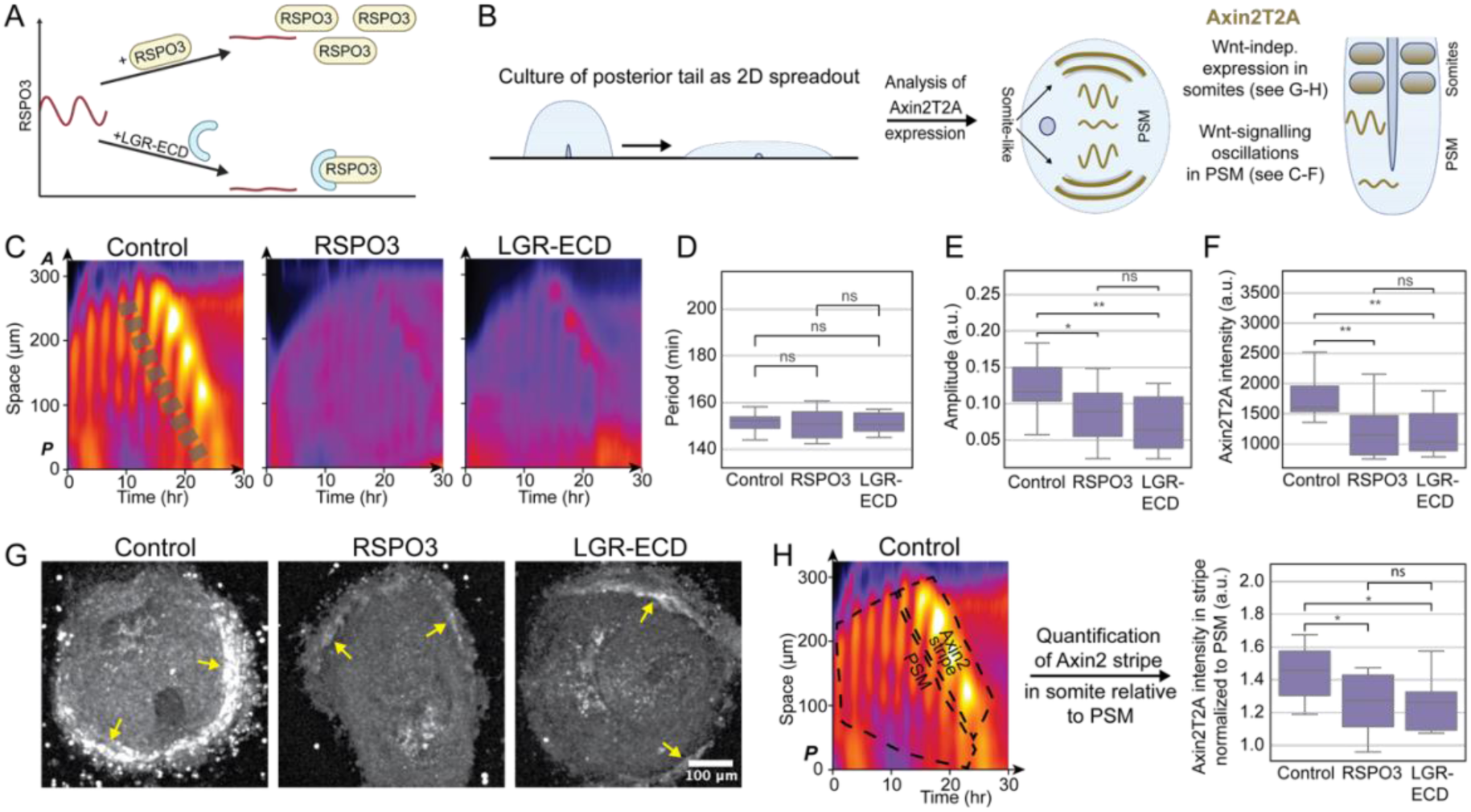
RSPO3 modulates Wnt-signalling oscillation amplitude and decreases anterior differentiation robustness. **A** Schematic summarizing the predicted effect of excess RSPO3 or blocking RSPO3 increasing or decreasing free extracellular RSPO3 levels. **B** Schematic representation of culturing E10.5 mouse PSM 2D spread-out cultures expressing Axin2T2A-Achilles. *Left* posterior tailbud explants spreading in a fibronectin-coated dish. *Centre* Wnt signalling-dependent Axin2T2A waves in 2D spread-out cultures travel radially from centre to periphery. This corresponds to Axin2T2A waves travelling from posterior to anterior in the tail (*right*). In periphery of 2D spread-out cultures and in somites of embryonic tails Axin2T2A expression is stable and not dependent on Wnt signalling. **C-H** E10.5 mouse PSM 2D spread-out cultures expressing AxinT2A-Achilles were incubated with 5 % RSPO3, 5 % LGR5-ECD or 5 % Control medium and fluorescence real-time live-cell imaging was performed (c_ontrol_ = 18, n_RSPO3_ = 12, n_LGR5-ECD_=9, N = 5). **C-F** Effect on Wnt-signalling oscillations in PSM: **C** Representative kymographs along a line from centre-to-periphery of spread-out cultures. **D-F.** Quantification of Axin2T2A timeseries data in anterior PSM (corresponding to the dashed line in **C**): oscillation period **D**, oscillation amplitude **E** and signal intensity **F**. **G-H** Effect on somite differentiation: **G** Representative image of Axin2T2A expression in spread-out cultures at the peak of peripheral Axin2T2A stripe formation. Yellow arrows: Axin2T2A stripe in somite-like structures. Scale bar 100 µm. **H.** Quantification of Axin2 stripe in forming somites. *Left panel:* Example kymograph (control in **C**) illustrating the ROIs used to quantify Axin2T2A intensity in forming somites. *Right panel*: Quantification of Axin2 stripe intensity relative to PSM intensity. In boxplots, each point represents the mean of one biological sample. For period **D**, amplitude **E**, statistical significance was assessed by one-way ANOVA followed by Tukey’s multiple comparisons test. For Axin2 intensity **F**, data did not meet the assumptions for normality (Shapiro–Wilk test), so statistical differences were tested by Kruskal–Wallis test followed by Dunn’s post-hoc test with Holm adjustment for multiple comparisons. *p* < 0.05 was considered significant. (* p<0.05, ** p< 0.01, ns non-significant).

Thus, our data indicate that interfering with dynamic RSPO3 expression prevents high-amplitude Wnt-signalling oscillations and affects downstream somite differentiation.

## Discussion

Here, we analysed spatial and spatiotemporal gene and protein expression in mouse somitogenesis, confirmed known factors and revealed many new potentially dynamic transcripts and proteins in the embryonic tail. Based on this approach, we uncovered a mechanism in which dynamic antagonistic ligand–receptor patterns modulate signalling oscillations during embryo development (Fig. 7).

**Figure 7.**
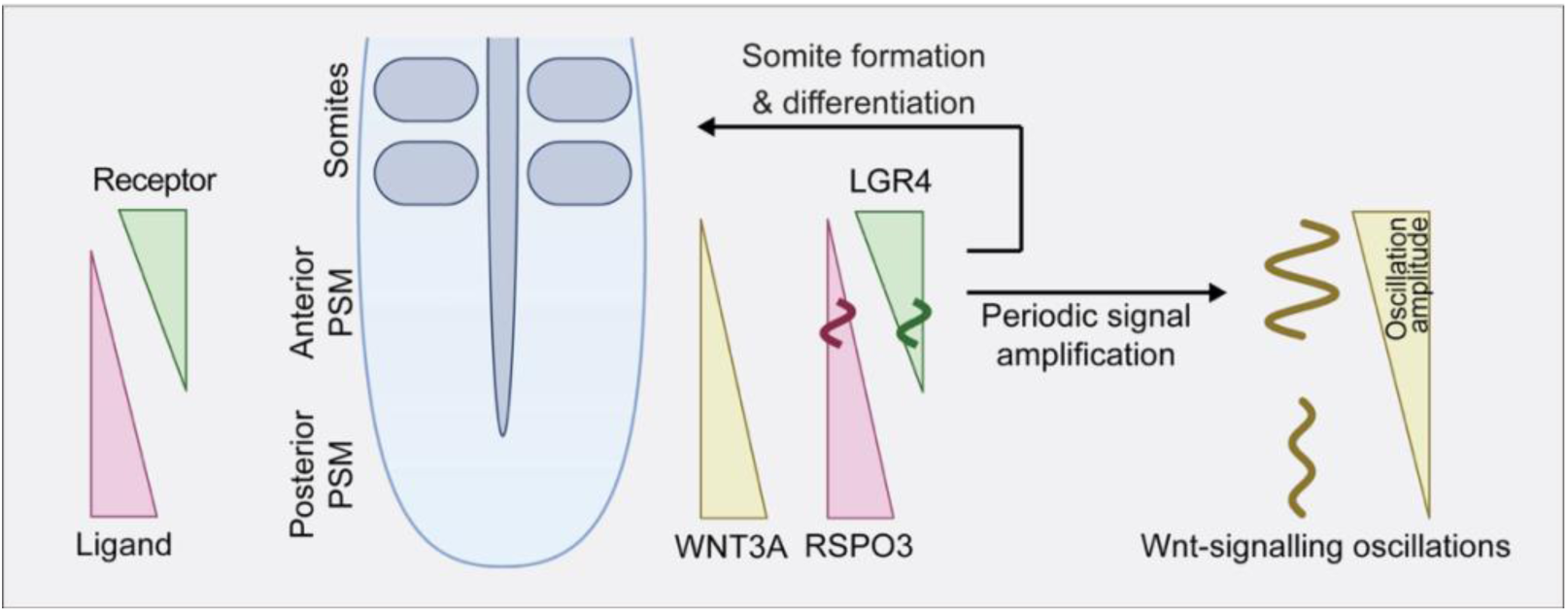
Dynamic antagonistic gradients of RSPO3-LGR4 regulate Wnt-signalling 863 oscillation amplitude and differentiation of anterior PSM. General antagonistic ligand-receptor expression pattern found with omics data (*left*). Proposed working model: Dynamic antagonistic gradients of RSPO3 and LGR4 in anterior PSM induce periodic signal amplification of Wnt signalling leading to high-amplitude oscillations and are required for somite differentiation.

We provide one of the first spatial proteome maps of somitogenesis. We identified posteriorly enriched proteins, including morphogens and signalling components, and proteins increasing in anterior PSM or somites, including FOXP1, a candidate regulator^34,59^ and somite marker. The cytoskeletal component EZRIN also showed graded expression with highest levels in posterior PSM (and overlying skin), potentially reflecting mechanical differences in the tissue consistent with fluid-to-solid transitions observed in zebrafish^60–62^. These data provide a foundation for exploring how cytoskeletal and membrane proteins shape tissue mechanics and morphogenesis during axial elongation.

When correlating mRNA and protein expression patterns to each other, we found that many mRNA–protein pairs showed corresponding spatial patterns, such as Wnt3a, supporting the long-standing use of *in situ* hybridization and transcriptomics as reliable proxies for protein localisation. However, numerous mRNA-protein pairs had low or inverse correlations, highlighting the need to study protein dynamics directly, as post-transcriptional mechanisms, differential stability and translational control can decouple RNA and protein levels (e.g.,^31,32,63,64^). Interestingly, many poorly correlated mRNA–protein pairs were associated with metabolic processes. Given the emerging regulatory role of metabolism in development^65–70^, further investigation of enzyme expression and metabolic flux in embryogenesis is warranted. Previous work largely focused on identifying cyclic genes in posterior PSM at the RNA level^26^. Our work extends this by including dynamic genes in anterior PSM and generating a list of candidate dynamic proteins. Using neural network–based analysis, many proteins were predicted to oscillate (p < 0.05). Although most did not pass multiple testing correction and may include false positives, several known cyclic transcripts and proteins showed similar significance levels, and we validated dynamic expression of selected candidates through staining of staged embryos or reporter assays. Combined, this highlights the validity of the selected significance thresholds. Conversely, due to data noise, it is likely that additional dynamic proteins were undetected. Time series data from these candidates may still provide valuable insights into their spatiotemporal expression patterns. Besides classic clock components, we identified dynamic candidates involved in metabolism, cell mechanics and centrosomal function. Whether these proteins contribute directly to the segmentation clock or reflect downstream responses^28^ remains an open question. Our dataset thus provides a rich resource for studying the interplay between oscillatory signalling, tissue dynamics and somite formation with spatial and temporal resolution.

Notably, absence of a protein in our data might reflect detection limits rather than true absence (as in the case of HES7^71^). Future advances in the sensitivity of mass spectrometers will enable deeper, spatially resolved insights into tissue regulators, cellular heterogeneity and post-translational modifications that might influence segmentation clock dynamics and somite development. Moreover, we noticed a higher variability in expression levels between the oscillation peak/trough in contrast to the peak or trough. This is likely due to more rapid changes in molecular levels during transitions, compared to the relatively stable concentrations at maxima and minima. Increasing sampling points across the oscillation cycle could help reducing this apparent noise. However, current methods require many embryos, raising ethical and logistical constraints. Future improvements in mass-spectrometry sensitivity may allow higher-resolution time series from fewer biological samples.

Based on our omics data, we addressed how morphogen gradients regulate downstream signalling. We focussed on the RSPO–LGR pathway, known to amplify Wnt signalling^46,50,51^. RSPO3 knockout mice are embryonically lethal around E10.5 due to placental failure and show delayed development with decreased trunk size^53^, reminiscent of the vestigial mutation showing decreased *Wnt3a* levels^72^, highlighting the essential role of RSPO3 in embryonic development. We found that both RSPO3 and its receptor LGR4 displayed dynamic expression in anterior PSM. When interfering with the dynamics of RSPO3 by either providing it in excess or blocking its function, amplitude and levels of Wnt-signalling oscillations decreased and somite differentiation was impaired. The dampening effect of excess RSPO3 on Wnt signalling might be due to negative feedback loops upon overactivation^73^. Importantly, it strongly suggests that the dynamics of the RSPO-LGR pathway are essential for regulating signalling oscillations in anterior PSM. In the noisy environment of the cell, decreasing the Wnt-oscillation amplitude by interfering with RSPO-LGR4 might lower the signal-to-noise ratio and decrease the robustness of differentiation and somitogenesis. Therefore, we propose a dynamic antagonistic ligand-receptor expression pattern as a mechanism to amplify signalling dynamics for robust differentiation and somite formation (Fig. 7).

We also found antagonistic ligand-receptor expression patterns for other signalling pathways including FGF and Notch, suggesting that similar mechanisms might modulate dynamics of other pathways within the PSM. Differential or antagonistic expression of signalling components including ligands and their receptors is being revealed in other systems, including Wnt signalling in the *Drosophila* wing disc or Bmp signalling in mouse embryos (e.g. ^74,75^). Dynamics in signalling components might also be important in these contexts. Together, our study highlights the necessity of considering dynamics of both signalling ligands and receptors and the resulting signalling activity when investigating morphogen gradients and tissue-level developmental processes.

Our dataset provides an invaluable resource for mechanistic studies of posterior embryonic development – including somitogenesis, axis elongation, tissue mechanics and metabolism – and offers insights relevant to congenital disorders affecting somite-derived structures and regenerative medicine. Similar integrative approaches hold promise for dissecting dynamic processes in other contexts such as tissue regeneration and homeostasis.

## Methods

### Animal use statement + study protocol

All experimental procedures were performed in accordance with institutional guidelines and approved by the Ethical Review Board of the Hubrecht Institute. The Animal Experimentation Committee (DEC) of the Royal Netherlands Academy of Arts and Sciences (KNAW) reviewed and approved all animal protocols under DEC project permits AVD801-0020186824 and AVD80100202316978. Mice were bred and housed according to internal guidelines at the Hubrecht animal facility, with ad libitum access to water and standard chow, in a controlled environment maintained at 50–60% humidity, 22–23°C temperature, and a 12-hour light/dark cycle.

### Mouse Strains

All transgenic mouse lines used in this study have been previously described. The Achilles-Hes7^34^ line was generously provided by Ryo Kageyama, the LuVeLu line^17^ by Olivier Pourquié and Axin-T2A-Venus line^18^ by Alexander Aulehla. We used C57BL/6N mice for microfluidic experiments intended for either RNA-seq or proteomics. Other wild-type mice used in this study were either C57BL/6N, C57BL/6J or F1 hybrids (C57BL/6N*C3H/N).

### Animal experiments

Female mice were euthanized at 10.5 days post-coitum (dpc), and embryos were dissected. Dissections were carried out in cold PBS, for omics methods, immunofluorescence or ISH/HCR experiments, or Embryo Culture Medium (ECM) (phenol-red free DMEM/F12 supplemented with 1% bovine serum albumin (Biowest, P6154), 200 μg/mL penicillin-streptomycin (Gibco, 15140122), 1× GlutaMAX (Gibco, 35050038), and 10% glucose solution (Sigma-Aldrich, G8769)) for whole mount and spread-out tail tip tissue culture in either culture dish or microfluidic chip. Tails intended for ISH, HCR or immunostaining were fixed overnight at 4 °C with 4% PFA. For ISH or HCR, tissue was subsequently dehydrated to 100% methanol and stored at –20 °C, while for immunofluorescence tails were stored in PBS+0.025%TritonX, and stored at 4 °C until use.

### Sectioning of embryonic tissue for immunofluorescence

Sectioning of embryonic tails for subsequent immunofluorescence was done as previously described^76^. Briefly, E10.5 mouse embryonic tails were placed in a plastic mold, embedded in 4% low melting agarose (LMA), and incubated for 15 min on ice to solidify the gel. The resulting LMA block was carefully removed from the mold and affixed to the specimen stage using cyanoacrylate glue. The stage was then submerged in a precooled phosphate-buffered saline (PBS) bath within the vibratome chamber, ensuring the LMA block was fully immersed by adding additional cold PBS as needed. Tissue sections approximately 70 μm thick were cut using a vibrating razor blade microtome set to consistent parameters across samples: cutting speed of 0.5 mm/s, amplitude of 0.9 mm, and vibration frequency of 65 Hz to minimize tissue damage. Sections were gently transferred from the PBS bath using a fine brush and microscope slide into a 6-well plate containing 2 ml PBS per well for subsequent processing.

Subsequent immunostainings were performed in the 6 well plate. Sections were permeabilized and blocked with PBS, 5% BSA, 1% Triton X-100 solution for 10 minutes at room temperature (RT), following primary antibody incubation overnight at 4°C in PBS, 2.5% BSA, 0.5% Triton X-100 with Ezrin (1:100 dilution) in the 6 well plates. Tissues were washed in PBS, 2.5% BSA, 0.5% Triton X-100 for 4x 30 min at RT. Conjugated secondary-antibody A647 (1:1000 dilution), Phalloidin 555 (1:500) and DAPI were incubated overnight at 4C in PBS, 2.5% BSA, 0.5% Triton X-100. Tissues were washed in PBS, 2.5% BSA, 0.5% Triton X-100 for 4x 30 min at RT and rinsed in PBS before imaging. Tissues sections were transferred to 35mm glass bottom dishes with minimal amount of PBS to avoid drying of the tissue while imaging.

### Embryonic tissue culture of whole tail and 2D explants

*Ex vivo* culture of samples was performed in ECM (as described above), according to previously described protocols^18^. In short, whole mount samples were cut between the 2^nd^ and 3^rd^ somite, while tails for spread-out tail tip tissue culture were cut before the neural tube. Both were subsequently cultured at 37 °C with 60%O2 and 5% CO2 in ECM, either in an incubator, or microscope with a gas- and temperature-controlled chamber. Small molecule inhibitors were dissolved in DMSO and used at the concentrations indicated in the main text and Fig. legends. These included CHIR99021 (Chiron; Sigma-Aldrich, SML1046), IWP-2 (Sigma-Aldrich, I0536), IWR-1 (Merck, I0161) and DAPT (Sigma-Aldrich, D5942). To control experiments, we added equal volumes of DMSO. 5% of RSPO3-containing medium, LGR5-ECD^49^ or Control were added to ECM. After culture, samples were fixed with 4% PFA in PBS for 30 min at RT, and either stored in PBS at 4C for immunofluorescent staining or dehydrated to 100% methanol and stored at –20°C for HCR.

### Generation of mESC RSPO3 reporter line

The RSPO3-T2A-Achilles knock-in reporter line was generated using CRISPR/Cas9 CRISPaint^77^ in IB10 ESCs. To generate RSPO3-T2A-Achilles, we targeted the stop codon of the endogenous RSPO3 locus. In agreement with the CRISPaint editing system we prepared 3 vectors: pSPgRNA (Addgene plasmid #47108) containing the RSPO3 sgRNA sequence (5’- AGTCAGCACTGTACACTAGA-3’); donor vector containing reporter cassette with the T2A-Achilles-NLS-PEST sequence and a selection cassette with Neomycin; and the corresponding Frame Selector pCAS9-mCherry-Frame +2. IB10 mESCs were co-transfected using Lipofectamine 3000 with 1.25ug of DNA per plasmid, selected for 1 week with 300 ug/ul of G418 and 100 clones were picked afterwards. Grown clones were genotyped and sequenced from RSPO3 C’-terminal region (Exon5) into the Achilles reporter.

### Gastruloid protocol

RSPO3-T2A-Achilles edited mESCs were maintained in 0.2% gelatinized 6 well plates with MEFS (prepared in house) with N2B27 media (DMEM/F12 without Phenol red, Neurobasal medium, B27 without Vit-A, GlutaMAX, Penicillin–Streptomycin) supplemented with 1x LIF, 3µM CHIR99021, 1µM PD0325091 and 20% KSR (20%), 0.1% β-Mercaptoethanol) in a humidified incubator (5% CO2, 37 °C) and passed every 2 days at a 1:16 or 1:20 ratio.

Gastruloids were prepared as previously described. In short, at day 0 cells were removed from MEFS: cells were washed with 1 mL of N2B27b (N2B27 supplemented with 0.1% β-Mercaptoethanol) and 200 ul of Accutase was added. After 3-5 minutes at RT, cells were collected and resuspended in 2 mL N2B27b and seeded in a 6 well plate and incubated for 1h (5% CO2, 37 °C). The media (containing mESCs) was collected to a 15 mL tube with additional 2 mL of N2B27b and centrifuged for 3.5 min at 0.2g. Cells were resuspended and additionally washed twice with 5 mL of N2B27b. After the last wash, cells were resuspended in 1 mL of N2B27b, the cell concentration was determined, and cells were diluted in N2B27b to a concentration of 7.5 cells per microlitre. 40 μl (with around 300 cells) of this suspension was transferred to individual wells of a U-bottomed 96-well plate (Greiner Bio-One, 650185). At 48h, gastruloids were supplemented with 150ul (per well) of N2B27b with 3 µM of CHIR99021, and at 72h were washed with N2B27b. At 96h gastruloids were washed with N2B27b, and the best ones (elongated gastruloids) were pooled in a 15 mL tube to be embedded in 10% Matrigel and N2B27b and transferred to a light sheet imaging chamber for live-cell imaging.

### Microfluidics

The full protocol for performing the microfluidics experiment has been described elsewhere^78^ with the following change: During bonding of the microfluidic chip to the glass slide the chip was flushed with PLL-PEG to prevent attachment of embryonic tissue. In short, the following steps must be performed:

#### PDMS Chip Fabrication

PDMS (SYLGARD 184, Dow Corning) was mixed at a 9:1 base-to-curing agent ratio, degassed in a desiccator and poured into microfluidic molds to a ∼500 µm thickness. The molds were degassed again to remove trapped air and cured overnight at 65 °C. Chips were cut out, holes punched (1 mm), cleaned with adhesive tape and bonded to glass slides using plasma treatment (1 min vacuum, 1 min gas to 0.37 mbar, 1 min plasma). Chips were immediately coated with PLL-PEG (1:20 dilution in HEPES, pH 7.4), incubated at 50 °C for 5 min, and at room temperature for 2 h.

#### Chip Preparation for Culture

Chips were flushed with PBS + 1% Pen/Strep and submerged in this solution to prevent air entry. Inlet/outlet tubing was UV-sterilized for 15 min. Outlet tubing was attached and PBS perfused using a syringe pump for ≥2 h or overnight.

#### Culture Medium and Drug Treatments

Syringes were loaded with culture medium (DMEM/F12 without Glucose and Glutamine, supplemented with 0.5 g BSA, 500 µL GlutaMAX, 55.5 µL 45% Glucose, 500 µL Pen/Strep, and 1.25 mL 1 M HEPES) or culture medium + drug (i.e., 2 µM DAPT + 10 µM Cascade Blue) or culture medium + control (DMSO + Cascade Blue) and degassed.

#### Embryo Tissue Loading and Setup

E10.5 mouse tail explants (LuVeLu, Axin2T2A-Venus or C57BL/6N tails) were dissected and loaded into chips using a P200 pipette. Remaining ports were sealed with PDMS plugs. Chips were connected to sterile tubing and placed in a humidified imaging chamber (37 °C, 5% CO₂, 60% O₂). Flow was stabilized at 20 µL/h for 20 min, then medium-only flow continued at 60 µL/h prior to imaging.

#### Live Imaging

Programmable pumps were used to alternatingly flush chips with culture medium or medium + drug/ control (100 min medium, 30 min drug pulses) to entrain Notch-signalling oscillations. Real-time imaging was performed with autofocus enabled. Cascade Blue fluorescence was used to monitor drug delivery. Flow was controlled to minimize shear stress and preserve tissue viability throughout the experiment. Embryonic tails were imaged using confocal imaging as described below using an N Plan 10x/0.25 objective with a resolution of 512 x 512 (pixel size 3.03 µm) and 8 z planes with 12 µm step size.

#### Entrainment + omics or HCR

For entrainment of wildtype tails, experiments were performed in standard incubator (37 °C, 5% CO₂, 60% O₂) and without the addition of Cascade Blue. At the end of the experiment, tails were harvested from microfluidic chip in ice-cold PBS using P200 pipette. For omics experiments, tails were manually dissected using Feather ophthalmology blade.

### TOPFlash assay

HEK 293 SuperTOPFlash (STF) cells, a Wnt reporter cell line, were cultured in G418 culture medium (GlutaMAX, 10% FCS, 1 % penicillin/streptomycin and 200 μg/mL G418 sulphate) and split 1:8 onto a 96 well cell culture microplate (Greiner, #655090) into 100 μl per well. Cells were cultured for 24 hours at 37 °C (5% CO2, 20% O2), after which 100 μl of the various dilutions of conditioned medium were added. All dilutions contained 10% Wnt conditioned medium, and 0.25% RSPO3 was added. These were again incubated at 37 °C (5% CO2, 20% O2) for 24 hours, after which 175 μl was pipetted off and 25 μl of Steady-Glo (Promega, E2510, lot 0000393038) was added. This was incubated for 20 minutes on a shaker at 300 rpm, after which luminescence was imaged using Berthold technologies Centro XS3 LB 960 with MikroWin 2010 (Labsis GmbH).

### Immunofluorescence

#### Mouse embryo tails (E10.5)

Tails were permeabilized and blocked with PBS, 5% BSA, 1% Triton X-100 solution for 2h at RT, following corresponding primary antibody incubation overnight at 4C in PBS, 2.5% BSA, 0.5% Triton X-100 in a rotating wheel at 9 rpm (rotations per min). Tissues were washed in PBS, 2.5% BSA, 0.5% Triton X-100 for 4x 30min at RT. Conjugated secondary-antibodies (listed in resource table were diluted at 1:500 or 1:1000) were incubated overnight at 4C in PBS, 2.5% BSA, 0.5% Triton X-100 with DAPI (Sigma, 5 μg/mL) to enable nuclear staining. Tissues were washed in PBS, 2.5% BSA, 0.5% Triton X-100 for 4x 30min at RT and rinsed in PBS before imaging. Individual tails were transferred to an 8 well Ibidi glass bottom chamber with PBS.

#### Gastruloids

At 120h the culture media was carefully aspirated from the light-sheet chamber and gastruloids were fixed with 4% PFA for 20 minutes at RT and washed with PBS. Gastruloids permeabilized with PBS, 5% BSA, 1% Triton X-100 solution for 10 minutes at RT. Blocking in PBS, 2.5% BSA, 0.5% Triton X-100 solution was done for 2 hours at RT. Subsequent steps of primary and secondary antibody incubations and washes were performed as described above for embryo tails, except the samples were kept at 4°C. Samples were then imaged in the light-sheet chamber containing PBS.

#### Antibodies

Primary antibodies used for immunostainings: Pax3 (Rabbit, 1:100), alpha-Tubulin (mouse, 1:1000), Foxp1 (Rabbit, 1:1000), Dll1 (Rat, 1:250), Ezrin (Rabbit, 1:100), Sox2 (Mouse, 1:2500), Tbx6 (Rabbit, gastruloids 1:500, tails 1:1000), T (Goat, 1:1000), Cdx2 (Mouse, 1:500).

### Hybridization Chain Reaction (HCR)

HCR buffers, amplifiers and probes for Rspo3, Lgr4 and Axin2 were supplied by Molecular Instruments. The HCR protocol was performed as published previously (^35^ and https://dx.doi.org/10.17504/protocols.io.7pyhmpw). To summarize, E10.5 tails were rehydrated, treated with proteinase K (1:2000 for 5 min at RT), and postfixed. Probes were hybridized to samples overnight at 37 °C (2pmol probe mixture). The next day, samples were washed with probe wash buffer and 5XSSCT, then amplified overnight in the dark at room temperature using amplification buffer and snap-cooled hairpins for the relevant amplifier sequence. Finally, samples were washed with 5XSSCT.

HCR probes for Hes7 and Pax3 were designed using the insitu_probe_generator^79^. The following parameters were used: max 5 A/T or G/C homopolymers length; probes were blasted against their cDNA and removed if highly matched; probes were limited to 5 probe pairs. The probe sequences can be found in Table 1.

**Table 1:**
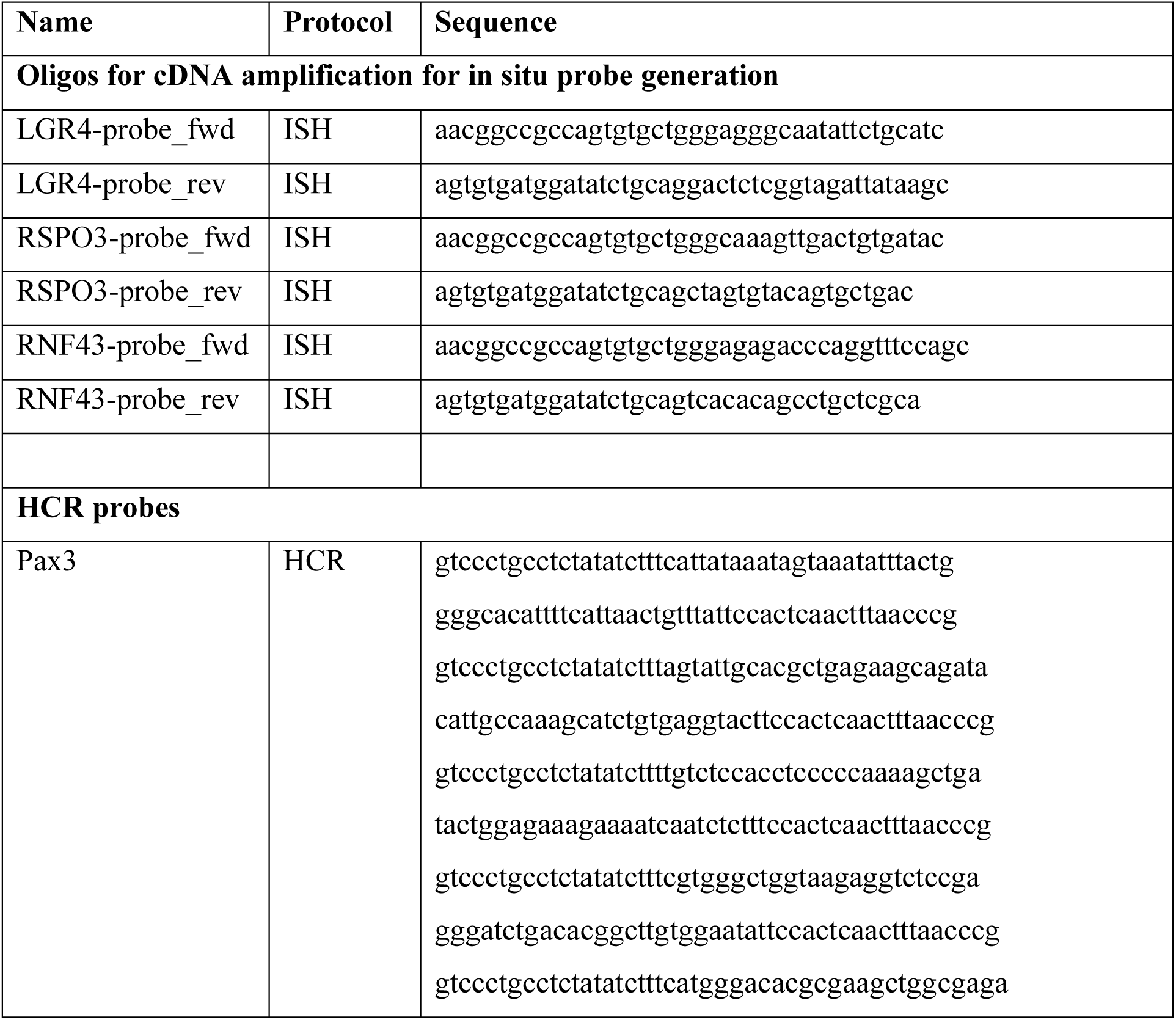
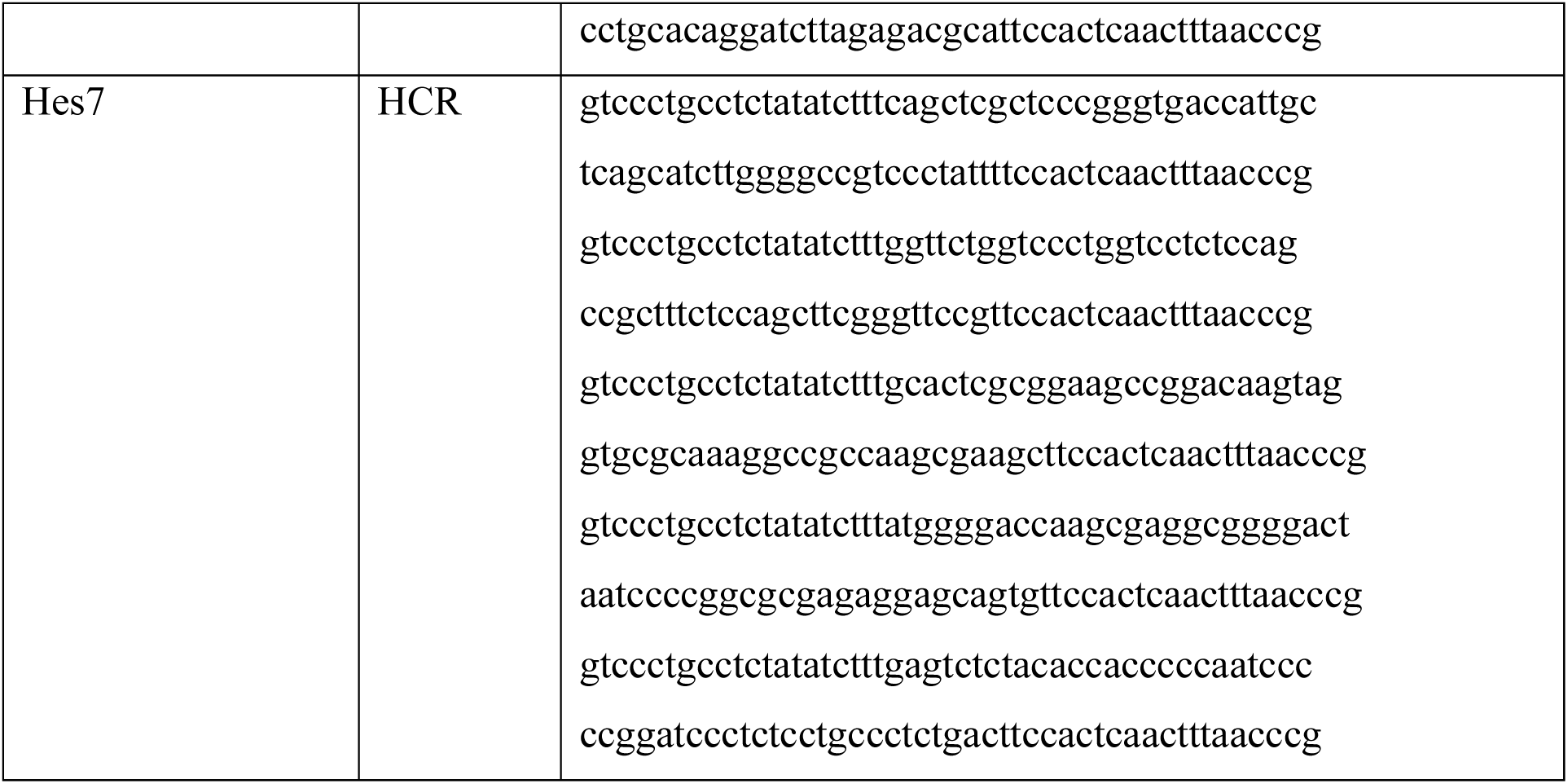
probes.

### ISH

Probe generation and *in situ* hybridization were performed as described previously (Aulehla et al., 2003; Lauschke et al., 2013). After rehydration, samples were bleached in 6% H2O2 for 5 min, then permeabilized by Proteinase K treatment (10ug/mL) for 12 min and postfixed in 4% PFA with 0.05 glutaraldehyde in PBS for 30 min. Probes were denatured at 80 °C for 10 minutes, after which they were hybridized to the samples using hybridization solution overnight at 68 °C. After washing, samples were blocked in TBST with 10% sheep serum for 2 hours, then incubated with anti-DIG AP antibody (1:1000) in TBST + sheep serum + 2% BSA at 4 °C overnight. Colour development was done with BCIP (170 ug/mL and NBT(340ug/mL) in NTMT at room temperature. *In situ* probes against Rspo1/2/3/4, Lgr4/5/6, RNF43 and ZNRF3 were generated using the primers shown in Table 1. *In situ* probe against Axin2 has been described previously^14^. Probes were synthesized using 10ug linearized plasmid with the Riboprobe Combination System (Promega). Images of in situ hybridizations were taken with a Leica MZ16F stereo microscope and a Leica DFC420C digital camera. Brightness and contrast were adjusted uniformly to the entire image.

### Imaging

#### Confocal live imaging

Confocal live imaging was carried out using a Leica TCS SP8 confocal microscope using a HC PL APO CS2 20×/0.75 dry objective for all imaging except for microfluidics experiments (see above). Images were acquired every 10 minutes using a z-step of 10 µm, covering a total depth of 100–150 µm with a z-resolution of 9 µm. Achilles or Venus fluorophores were excited using OPSL lasers at 514 nm. Live cell imaging was performed with resolution of 512 × 512 pixels, with a pixel size of 1.517 µm. For wholemount tissue imaging, samples were maintained in a humidified chamber at 37°C with 5% CO₂ and 60% O₂. For 2D cultures, the oxygen concentration was reduced to 20%.

#### Light-sheet Live Imaging

Live cell imaging of whole-mount embryo tails for Achilles-Hes7 phase analysis or gastruloids was performed using a Viventis LS1 Live light sheet microscope (Viventis Microscopy). Imaging took place in a humidified chamber at 37°C, with 5% CO₂ at either 60% O₂ (for whole tails) or 20% O₂ (for gastruloids). To visualize the Achilles fluorophore a 514nm laser was used. A 25× objective and 3.3 μm laser beam were used to acquire z-stacks of 60–80 steps, with 3 μm intervals. Images were captured at 2048 × 2048 pixel resolution every 15 minutes.

To stabilize whole-mount tissues while permitting normal growth, samples were embedded in 2% agarose in PBS, cast around a custom-designed insert within the sample holder. This insert contained five slots (1.5 mm × 0.5 mm) to hold individual tissues. For gastruloids, imaging was performed in a 4-well light sheet imaging chamber.

#### Light sheet fixed imaging of gastruloids

Immunostained gastruloids were imaged in the same light sheet chamber using the same parameters as for live cell imaging. In addition, capturing Phalloidin-405, RSPO3-T2A-Achilles and secondary antibodies A568 and A647 was achieved using lasers at 405 nm, 514nm, 564 nm and 647 nm wavelengths.

#### Confocal imaging of fixed tissue

To image immunofluorescent stainings or HCR, z-stacks of 10–12 planes were captured, with z-step sizes between 5-10 μm. Images were acquired at a resolution of 1024 × 1024 pixels (pixel size: 0.758 μm) using a Leica TCS SP8 MP confocal microscope equipped with an HC PL APO CS2 20×/0.75 or 10x dry objective. Excitation was achieved using OPSL lasers at wavelengths 365 nm, 488 nm, 546nm, 568 nm and 647 nm.

### Tissue preparation for spatial omics

After initial dissection, embryonic tails were dissected further into posterior PSM (p-PSM), anterior PSM (a-PSM) and the 2 last formed somites. For each region, tissue pieces from 8 embryos were pooled to ensure adequate sequencing/proteome depth. After pooling, excess liquid was removed using a 30G needle. For RNA-seq, 100uL TRIzol was added. For proteomics, samples were snap frozen in liquid nitrogen. Samples were then stored at −80°C until processing. 3 biological replicates were obtained from independent dissections for both RNA-seq and proteome analysis.

### Tissue preparation for spatiotemporal omics

After entrainment using microfluidics (see above), tissue samples were obtained from 4 consecutive timepoints of one oscillation cycle after 6 external drug pulses. To this end, embryonic tissue was recovered from the microfluidic chip and dissected in ice-cold PBS into p-PSM, a-PSM and the last formed somites. For each region, tissue pieces from 4 embryos were pooled to ensure adequate sequencing/proteome depth. After pooling, access liquid was removed using a 30G needle. For RNA-seq, 100uL TRIzol was added. For proteomics, samples were snap frozen in liquid nitrogen. Samples were then stored at −80°C until processing. For RNA-seq, 3 biological replicates from independent experiments were obtained. For proteomics, 4 biological replicates from independent experiments were obtained.

### Bulk-RNA sequencing

Bulk RNA sequencing was performed by Single Cell Discoveries (Netherlands) using a modified CEL-Seq protocol. Total RNA was extracted with the standard TRIzol reagent (Invitrogen) and used for library preparation. mRNA processing followed adapted versions of the single-cell mRNA sequencing protocol described previously^80,81^. Sample-specific barcodes were introduced during reverse transcription using CEL-Seq primers^82^, and samples were pooled after second-strand synthesis. The resulting cDNA was amplified by overnight in vitro transcription, after which sequencing libraries were prepared using Illumina TruSeq small RNA primers. Libraries were sequenced on the Illumina NextSeq500 platform in paired-end mode (read 1: 26 cycles; index read: 6 cycles; read 2: 60 cycles). Reads that mapped equally to multiple locations were discarded. Mapping and generation of count tables was done using the STARSolo 2.7.10b aligner.

RNA data have been deposited at the NCBI GEO repository (https://www.ncbi.nlm.nih.gov/geo/) with the identifiers GSE304260 and GSE304261.

### Mass-spectrometry

#### SP3 sample preparation

Samples were prepared using the SP3 method for low-input proteomics^83^. In short, cell pellets were lysed in Urea buffer (8M urea, 100mM Tris-HCl pH8.0, 10 mM DTT) and sonicated (10 cycles 30sec on/off) with a Bioruptor® Pico (Diagenode) followed by shaking at room temperature for 30 minutes. Then iodoacetamide (Sigma, cat. no. I1149) was added at a final concentration of 50 mM and incubated for 20 minutes at room temperature. For the SP3 bead method Sera-Mag™ SpeedBeads (Sigma, GE65152105050250) and Sera-Mag™ Carboxylate-Modified Magnetic Particles (Sigma, GE45152105050250) were mixed at a ratio of 1:1. Prewashed SP3 beads were added to the lysate followed by the addition of 10 volumes of acetone. Samples were incubated at room temperature for 20 minutes. Using a magnetic rack, the beads were immobilized and washed with acetone (Sigma, 270725), twice with 80% ethanol and once with acetonitrile (Biosolve, 1204101BS). Airdried beads were resuspended in 25ul of 0.1M TEAB (Sigma, T7408) plus 0.1ug sequencing grade trypsin (Promega, V5280). Proteins were digested for two hours while shaking at 37°C, after which the supernatants were collected. TEAB (0.1M) was added to the beads, incubated in a thermoshaker for 5min and supernatants were combined with the first elution. Digestion was continued overnight at 37°C. The next day, peptides were acidified by adding 10 μl of 10% v/v TFA (Biosolve, 202341) and purified on C18 Stagetips^84^.

#### Measurement

Samples were measured using a 60-minute reverse phase gradient of buffer B (80% acetonitrile, 0.1% formic acid) on an Easy-nLC1000 coupled on-line to an Orbitrap Exploris 480 (Thermo Fisher Scientific) operated in Top20 mode with a dynamic exclusion list of 45 seconds. The full scan of the peptides was set to a resolution of 120,000 in a scan range of 350–1300m/z. The normalized AGC target was 300% and the maximum injection time was set to 20 ms.

#### Label-free quantification

Raw files were analysed using Proteome Discoverer software (version 3.0) with Sequest HT and Chimerys™ search algorithms. Mouse Fasta database updated in 2017 from Uniprot was used as a search database. Protein group files were imported into R for downstream analysis using the DEP R package. Proteins identified with fewer than two peptides and those not annotated as “IsMasterProtein” were filtered out. For spatial data, proteins quantified in all triplicate samples of at least one sample group were considered for downstream analysis. For spatial data, missing LFQ values were imputed for statistical analysis using random draws from a manually defined left-shifted Gaussian distribution (for MNAR). Significant expression was defined as p-value < 0.05 and absolute lfc > 1. Spearman correlation between RNA and protein expression were calculated based on publicly available code^85^. For spatiotemporal data, missing values were not imputed. Further analysis and detection of dynamic proteins is described below. Mass-spectrometry data have been deposited at the ProteomeXchange Consortium through the PRIDE partner repository (https://www.ebi.ac.uk/pride/) with the identifier PXD064068.

### RNA-seq analysis

Bulk RNA-seq data was analysing using the python package PyDeSEQ2 using standard parameters. Genes with counts below 10 were excluded. Normalized counts were used for subsequent analyses.

### Omics data analysis

#### Gene ontology analysis

Gene ontology analysis was performed using enrichGO and GSEA functions of R package clusterProfiler^86^ (4.2.2)^86^.

#### Identification of dynamic genes

Identification of dynamic genes and proteins was performed using BIO_CYCLE^43,87^, a deep learning approach. Code was retrieved from https://circadiomics.ics.uci.edu/biocycle. Analysis was performed on non-imputed data. Datasets were filtered for genes that included at least one value per timepoint before BIO_CYCLE analysis. The neural network was retrained on posterior data from either RNA-seq or Proteome data setting the genes Lfng, Hes7, Dusp4, Dusp6, Axin1 and Axin2, if present in the dataset, as GroundTruth to create 2 separate trained models. Subsequently, RNA-seq and Proteome data for each region was analysed using the respective model setting 110 and 150 min as upper and lower period limits, respectively. on. Before further analysis, bulk RNA-sequencing data were subsequently filtered for genes, for which each timepoint contained at least 2 non-zero/NaN values. Mass-spectrometry data was filtered for proteins, for which each timepoint contained at least 2 non-zero/NaN values AND for which no more than 6 values were missing in total.

### Analysis of tail stainings

#### HCR

Tail images were quantified in Fiji by drawing a 50pixel-width, segmented line following the curve of the tail and encompassing the entire tail. Display range was adjusted to visualize control tails and subsequently adapted to all other images.

#### Immunostainings

In Fiji, z-slices were subjected to a maximum intensity projection (MIP) and background was subtracted from all images analysed. For quantification of Foxp1, Ezrin, Tubulin and phalloidin a polygon line was hand drawn in the regions corresponding to p-PSM, a-PSM and somite. The mean intensity for each region was plotted in boxplots using a python in a custom made Jupyter notebook.

For Pax3 stainings, in Fiji, acquired z-slices (5-7 µm per tail) were subjected to MIP, and a user defined segmented line (width varying 20 – 100 pixels) was drawn in each tail from posterior until somite 2 (as shown in corresponding representative image). The display range is the same for corresponding images shown. Extracted values from segmented lines were post processed in python in a custom made Jupyter notebook. Pax3 intensities were smoothed using gaussian filter (σ = 5) and tail distances in µm were averaged using rolling average (window = 9). The data was then binned (bins = 300) using the tail distance from posterior and normalized from 0-1.

## Data analysis

### Generation and analysis of kymographs

Analysis of oscillations in embryonic tails in microfluidics entrainment experiments were done the following way: Using Fiji (Schindelin et al., 2012), z-stacks were projected using a maximum intensity projection and time series were further processed with a Gaussian filter (5 px). Kymographs were created along a user-defined line from posterior to anterior (shown in corresponding representative images) of 2D PSM cultures or gastruloids using the plugin KymoResliceWide (intensity averaged over the width (20-200 px) of a wide line) available in Fiji.

To analyse oscillations in 2D mPSM cultures, z-stacks were first projected using maximum intensity projection. Time series were then smoothed using a Gaussian filter (10 µm). Kymographs were generated along a user-defined posterior-to-anterior axis and further smoothed with a Gaussian filter (σ = 10 µm).

Phase kymographs were produced as described previously^45^. Wave speeds were determined from the slope of lines in phase kymographs. Wave slopes were quantified by manually selecting the posterior onset of each wave in the phase kymograph; for each wave, phases between 0.6π and 1π were used to automatically compute a regression line in MATLAB^18^.

### Analysis of entrainment experiments

Experiments were analysed as described previously^18^. Kymographs were generated using the *KymoResliceWide* tool as described above along a 100-pixel-wide line drawn through the posterior-to-anterior axis of the sample. Signal intensity was then quantified within the kymograph by extracting mean intensities along 25-pixel-wide lines in the posterior and anterior PSM regions.

To combine experiments, the resulting time series data from individual experiments were manually aligned based on the timing of drug pulses to allow comparison across experiments. Data were smoothed using a moving average with a window size of 5 (corresponding to 50 minutes, with one image taken every 10 minutes). To remove slow trends, the data were subsequently detrended by subtracting a second moving average with a window size of 13. The aligned and detrended time series from individual samples were plotted together to assess synchronization across experiments.

Somite size was quantified using brightfield imaging by manually measuring both the length of the PSM and the length of newly formed somites over the course of the movie using Fiji^88^. Each measurement was performed in triplicate and averaged. Somite size was then analysed relative to the total PSM length to account for size variability between samples.

### Analysis of Axin2T2A signal in 2D explant cultures

Kymographs were generated as described above. Signal intensity was then quantified within the kymograph by extracting mean intensities along 10-pixel-wide lines in anterior PSM regions next to the anterior Axin2 stripe (see Fig. 6). We analysed Axin2T2A reporter signals using wavelet-based time-series analysis using the *pyBOAT* Python package^89,90^. The sampling interval was set to 10 minutes, and the period search range was defined as 50–300 minutes (100 linearly spaced values). To remove slow trends, raw Axin2T2A intensity signals were smoothed using sinc filtering with a cutoff period (Tₚ) of 210 minutes. The resulting trend was subtracted from the raw signal to yield a detrended time series.

Wavelet power spectra were computed on the detrended data, and the dominant oscillatory component was extracted using ridge detection (power threshold = 0, ridge smoothing window = 10). Amplitude normalization was performed using a moving window of 150 minutes to account for baseline fluctuations. From the resulting wavelet ridge, the phase, amplitude, and instantaneous period were extracted for each signal.

Only data points within the time window of 300–1200 minutes and with wavelet power > 3 were used for downstream period and amplitude analyses. For intensity analysis, raw intensity values were included over the same time window regardless of wavelet power. Period, amplitude and intensity were averaged per sample before plotting.

### Analysis of reporter dynamics in gastruloids

In Fiji^88^, z-slices were subjected to a MIP and time series were further processed with a Gaussian filter (5 µm). A user defined round ROI in the gastruloids’ RSPO3-T2A-Achilles positive region (as shown in Fig. 5J) was used to detect raw signal for each time point to be subsequently analysed in pyBOAT to quantify signal oscillations and the oscillation period using wavelet analyses.

Raw fluorescence intensity time series were smoothed using a centred rolling average (window = 5 frames, equivalent to 75 minutes at 15-minute intervals). For phase-based analysis, signals were detrended using a sinc filter with a cutoff of 210 minutes (via the pyBOAT package^90^) and normalized using a sliding window of 150 minutes. Period values were extracted from the wavelet ridge of the normalized detrended signal.

### Fourier Transform and Dominant Period Extraction

For each individual signal (identified by a unique combination of position and experiment), we computed the Fast Fourier Transform (FFT). Signals were zero-centred, and frequency components were converted to periods using the sampling interval of 15 minutes. In cases where the time series spanned 1065 minutes, the first and second harmonics were excluded to avoid artifacts from long-term drift. The dominant period, corresponding to the peak in the amplitude spectrum, was identified for each track, and a kernel density estimate (KDE) was plotted across all tracks to visualize the distribution of dominant oscillatory periods.

### Average Power Spectrum

To analyse the overall frequency content across the entire dataset, each time series was padded or truncated to a uniform length of 72 time points (18 hours). After mean subtraction, the power spectrum was computed from the FFT of each signal, and the average power spectrum was calculated across all tracks. The resulting average power values were converted to periods and smoothed using a Gaussian filter (σ = 2). The average power spectrum was then visualized as a weighted histogram of periods, providing a population-level view of dominant oscillatory modes.

### Data visualization and statistical analysis

Experiments were conducted with a minimum of three independent replicates. Data was analysed and visualized using python and standard python packages such as matplotlib, seaborn, pandas, numpy, scipy and scanpy, and functions such as heatmap, boxplot, lineplot. Code is available upon request. Data can be accessed and visualized on our website (integrative thingy)

Tukey-style boxplots in which the box delineates the 25th and 75th percentiles, and the bar the median and the whiskers extend to 1.5 times the interquartile range (IQR). For each sample, the mean value was calculated and used for statistical testing. For direct comparisons between two conditions, unpaired T Test was performed. For the analysis of Axin2T2A signal and signalling dynamics, Data normality was assessed by the Shapiro–Wilk test and homogeneity of variances by Levene’s test. If data met assumptions of normality and equal variance, one-way ANOVA followed by Tukey’s multiple comparisons test was performed. Otherwise, the non-parametric Kruskal–Wallis test was used, followed by Dunn’s post-hoc test with Holm correction. Statistical significance was defined as *p* < 0.05.

### Resource Table

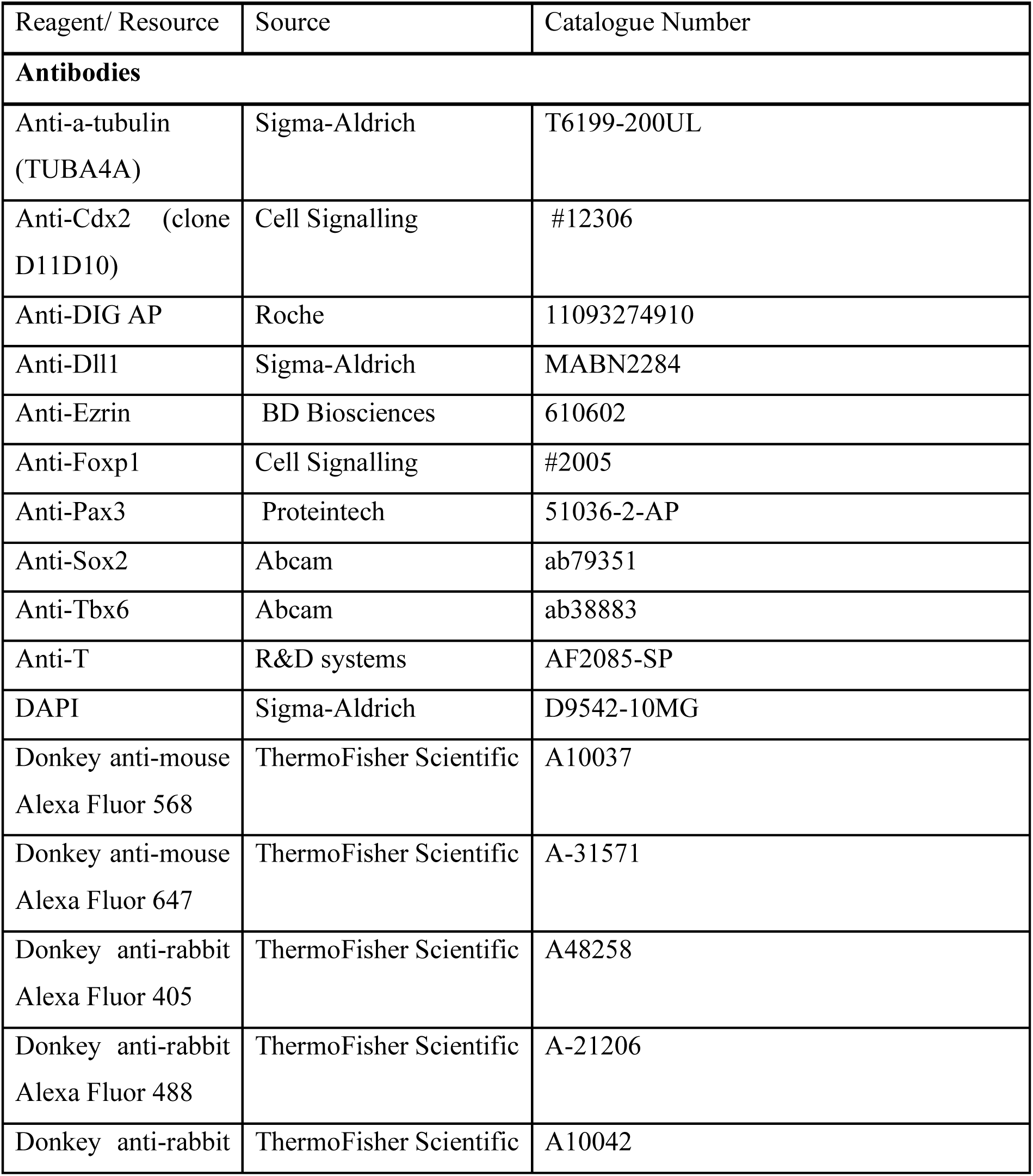
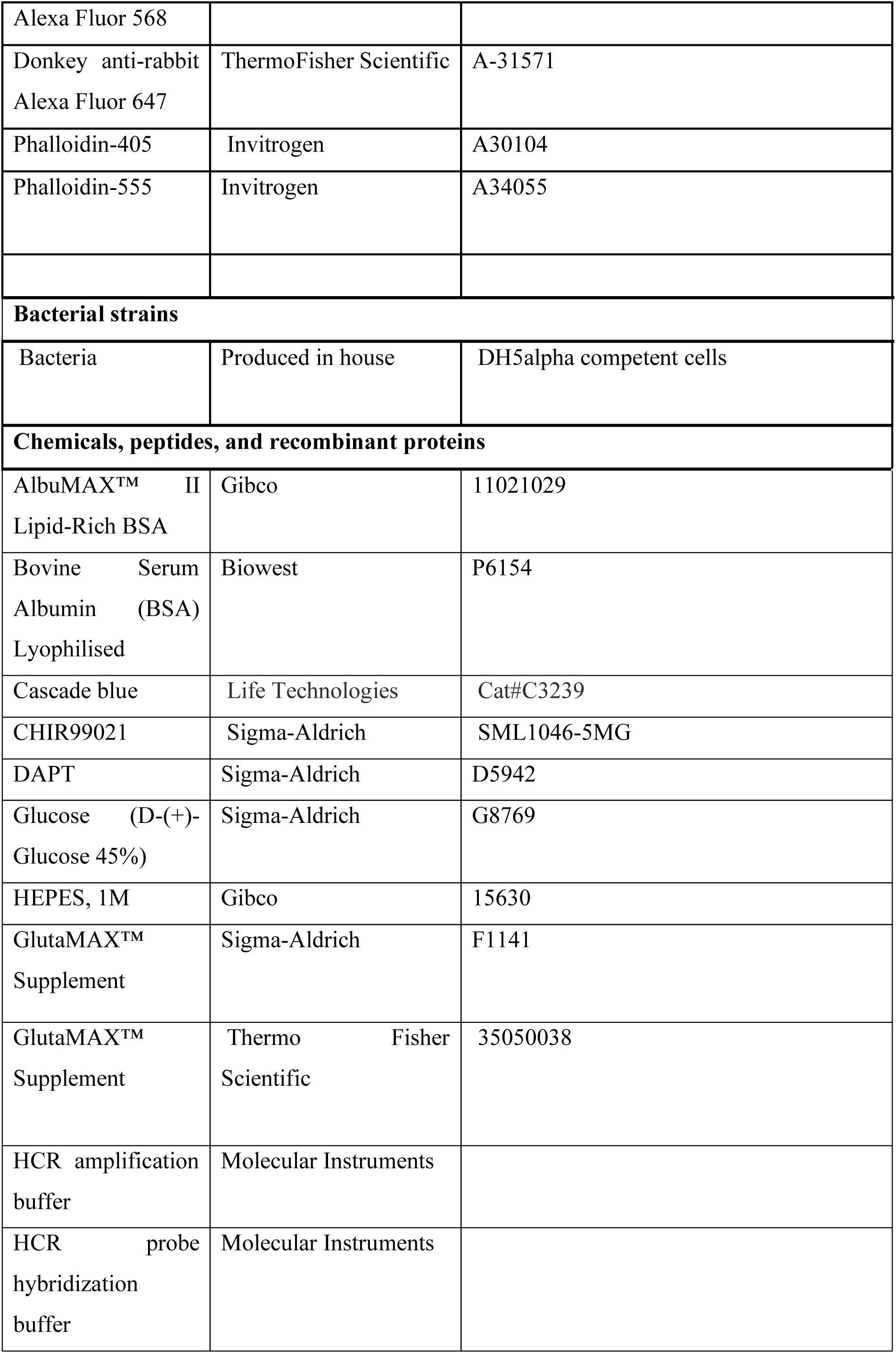
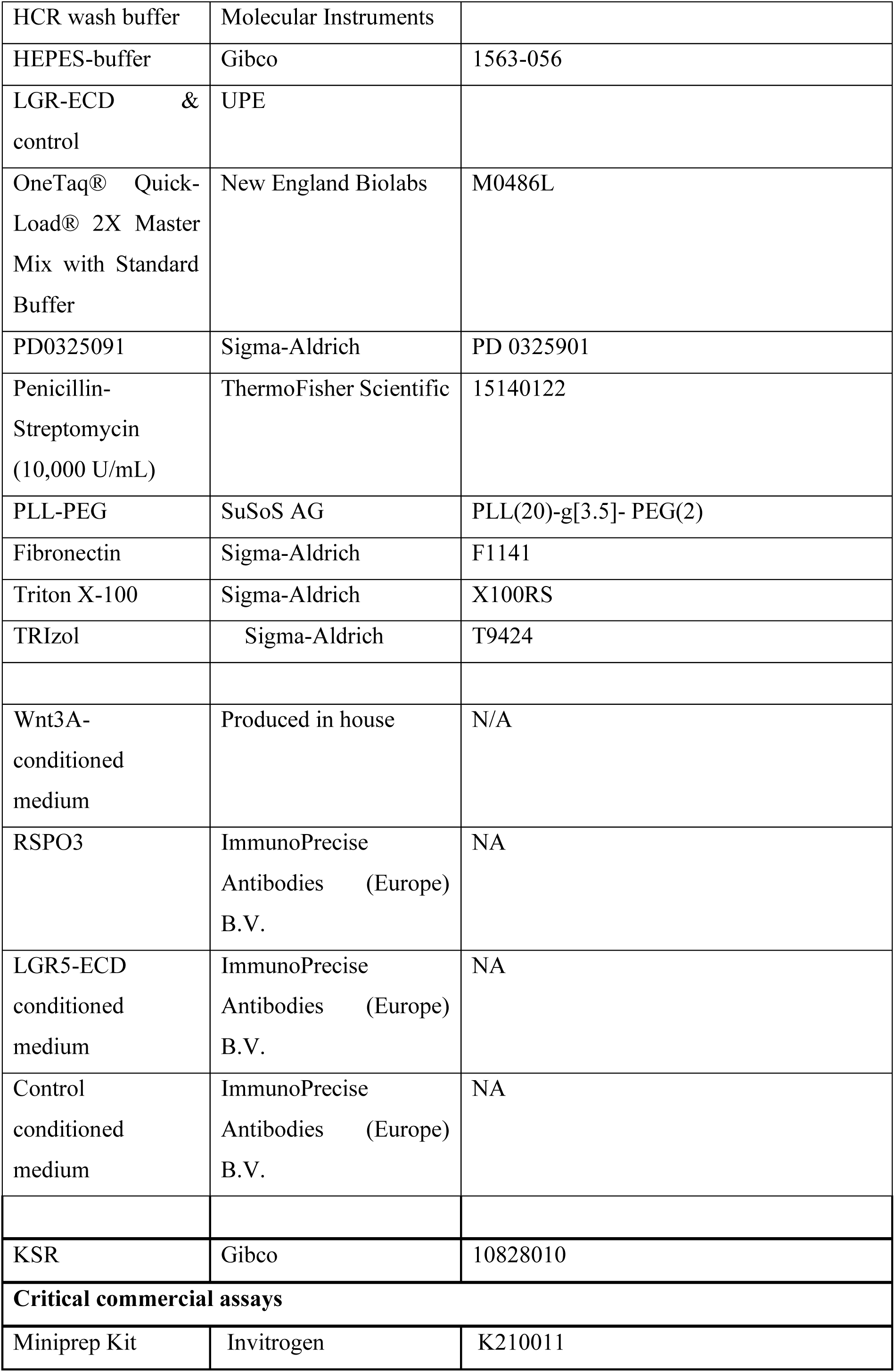
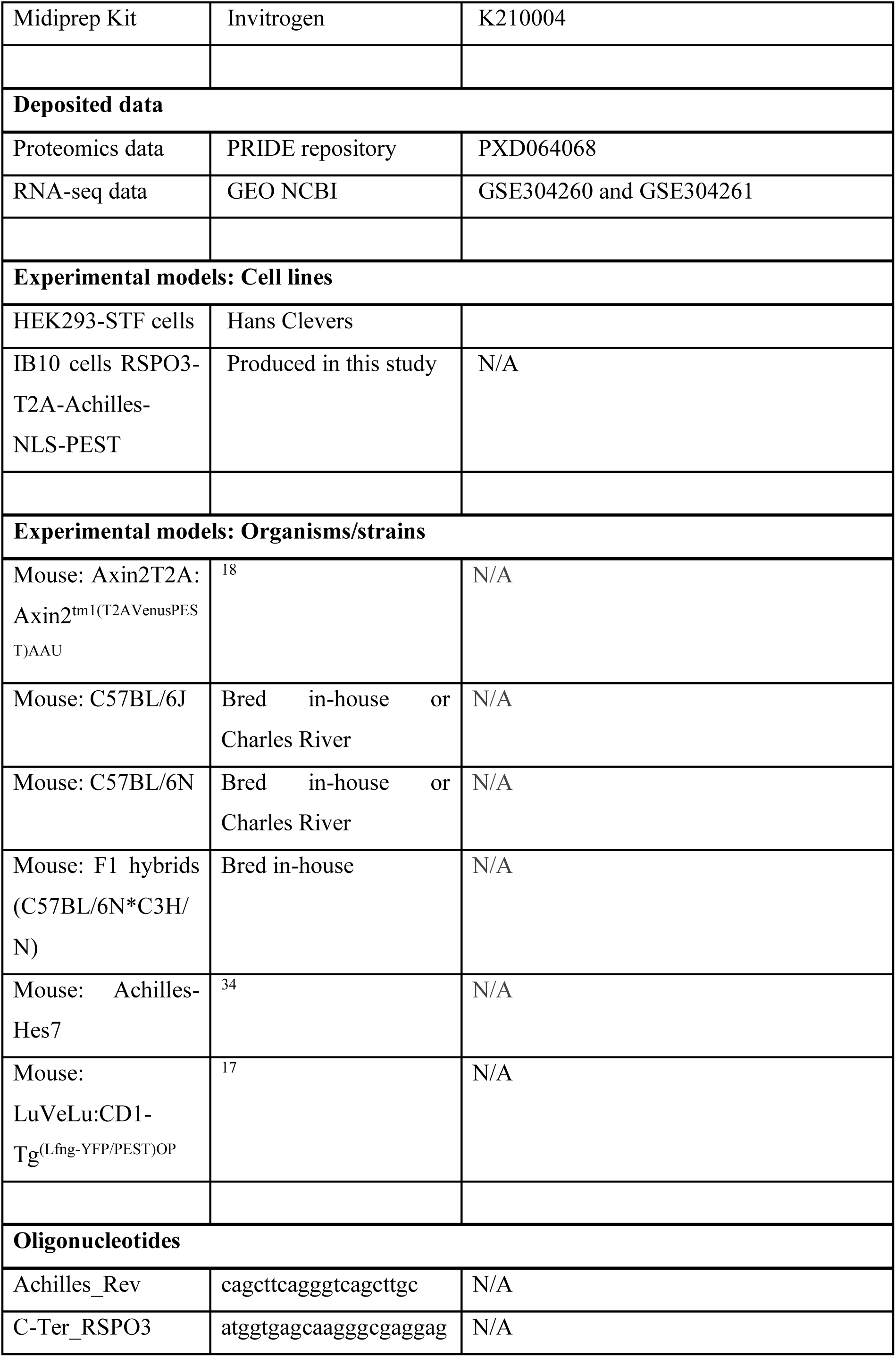
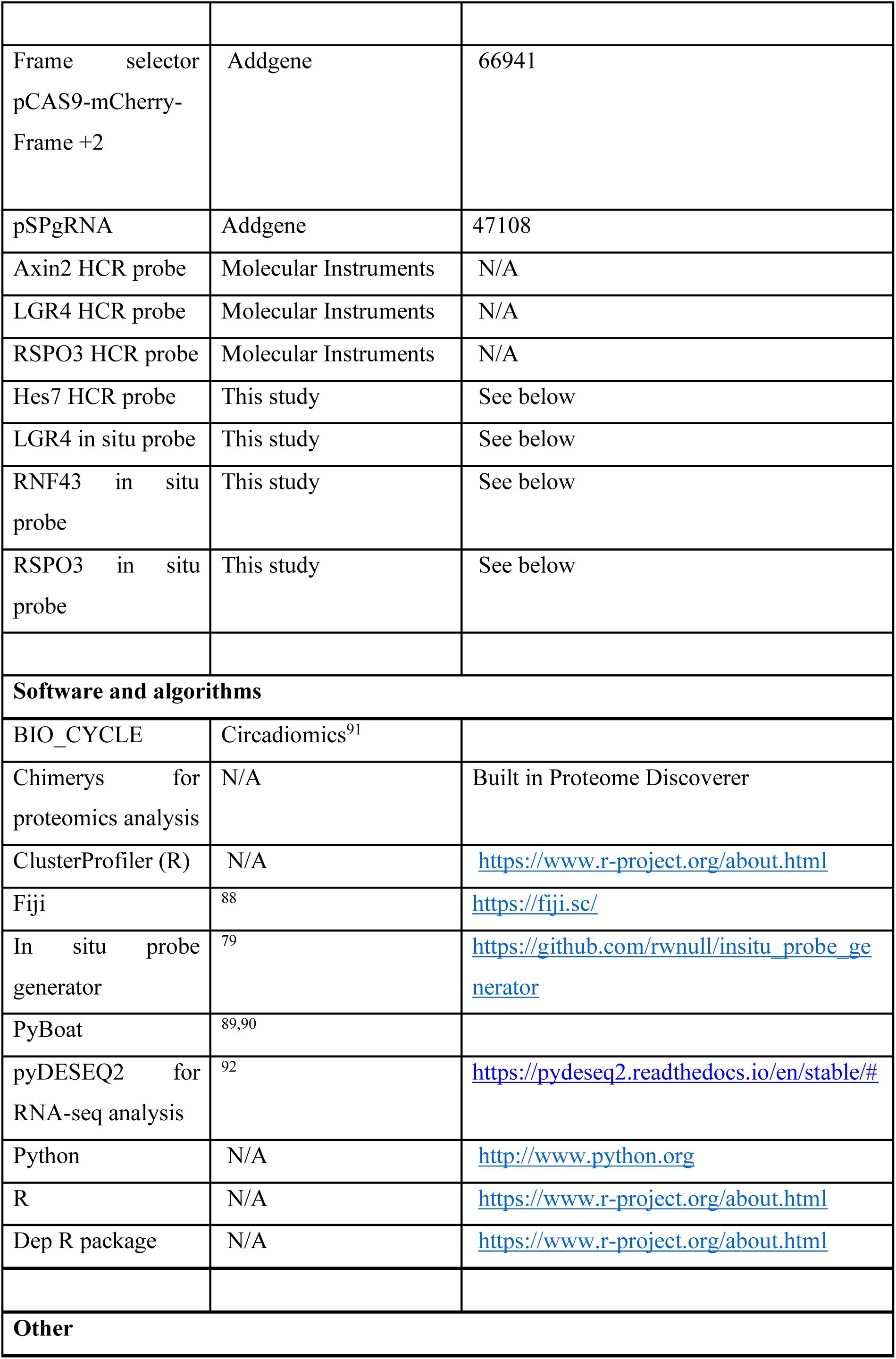
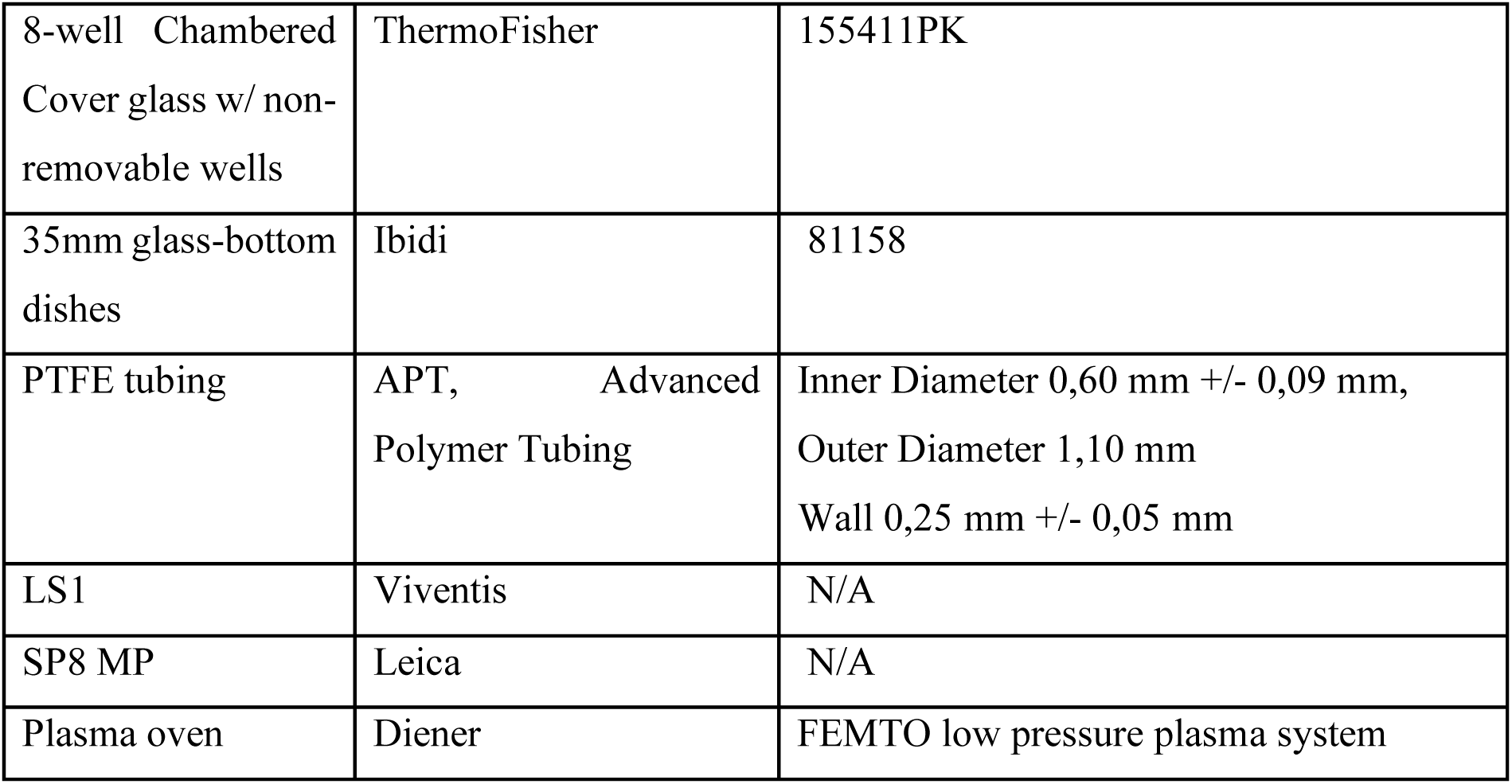

### Data and code availability

For requests contact the corresponding author. Mass-spectrometry data have been deposited at the ProteomeXchange Consortium through the PRIDE partner repository (https://www.ebi.ac.uk/pride/) with the identifier PXD064068. RNA data have been deposited at the NCBI GEO repository (https://www.ncbi.nlm.nih.gov/geo/) with the identifiers GSE304260 and GSE304261.

## Acknowledgments

We thank members of the Sonnen group, Anniek Stokkermans, Wim de Lau and Euan Joly-Smith for discussions and feedback and Alexander Aulehla for support. We thank the animal, FACS and imaging facilities of the Hubrecht Institute for their support. We are grateful to Nelleke Spruijt for help with mass-spectrometry and Jeroen Korving for help with tissue sectioning. We thank Pascal Jansen for technical support with mass-spectrometry. The LuVeLu reporter line was kindly provided by Olivier Pourquié, the Axin2T2A line by Alexander Aulehla, the Achilles-Hes7 by Ryoichiro Kageyama and the Achilles construct by Atsushi Miyawaki (Riken Center for Brain Science). The HEK293T Super TOPFlash reporter cell line and LGR5-ECD construct were kindly provided by Hans Clevers. This work was supported by the Hubrecht Institute and received funding from the European Research Council under an ERC starting grant agreement no. 850554 to and NWO Aspasia and M grants from the Dutch Science Organization to K.F.S. It also received funding from the ZonMW PSIDER grant 2021/15042. V.A. was supported with an EMBO postdoctoral fellowship. S.S. was supported with a VENI grant from the Netherlands Organisation for Scientific Research (NWO, VI.Veni.212.076). In addition, this research was supported by discussions of K.F.S. at Kavli Institute for Theoretical Physics (KITP), Santa Barbara, funded in part by grant NSF PHY-1748958.

## Author contributions

Conceptualization: W.H.M.M., V.A., S.S., M.V., K.F.S.; Methodology: K.F.S.; Validation: W.H.M.M., V.A., K.F.S.; Formal analysis: W.H.M.M., V.A., S.S.; Investigation: W.H.M.M., V.A., S.S., K.F.S.; Resources: W.M.T., M.J.O., E.I., C.G.S., K.P.; Data Curation: W.H.M.M., S.S.; Writing – Original Draft: W.H.M.M., V.A., K.F.S.; Writing – Review and Editing: S.S., M.J.O., M.V.; Visualization: W.H.M.M., V.A., S.S.; Supervision: M.V., K.F.S.; Project management: K.F.S.; Funding acquisition: V.A., S.S., M.V., K.F.S.

## Additional Information

Correspondence and requests for materials should be addressed to.

