## Extended Data for "Spatiotemporal proteomics reveals dynamic antagonistic gradients shaping signalling waves"

Extended Data Figures 1 - 7

### Extended Data Figure Legends

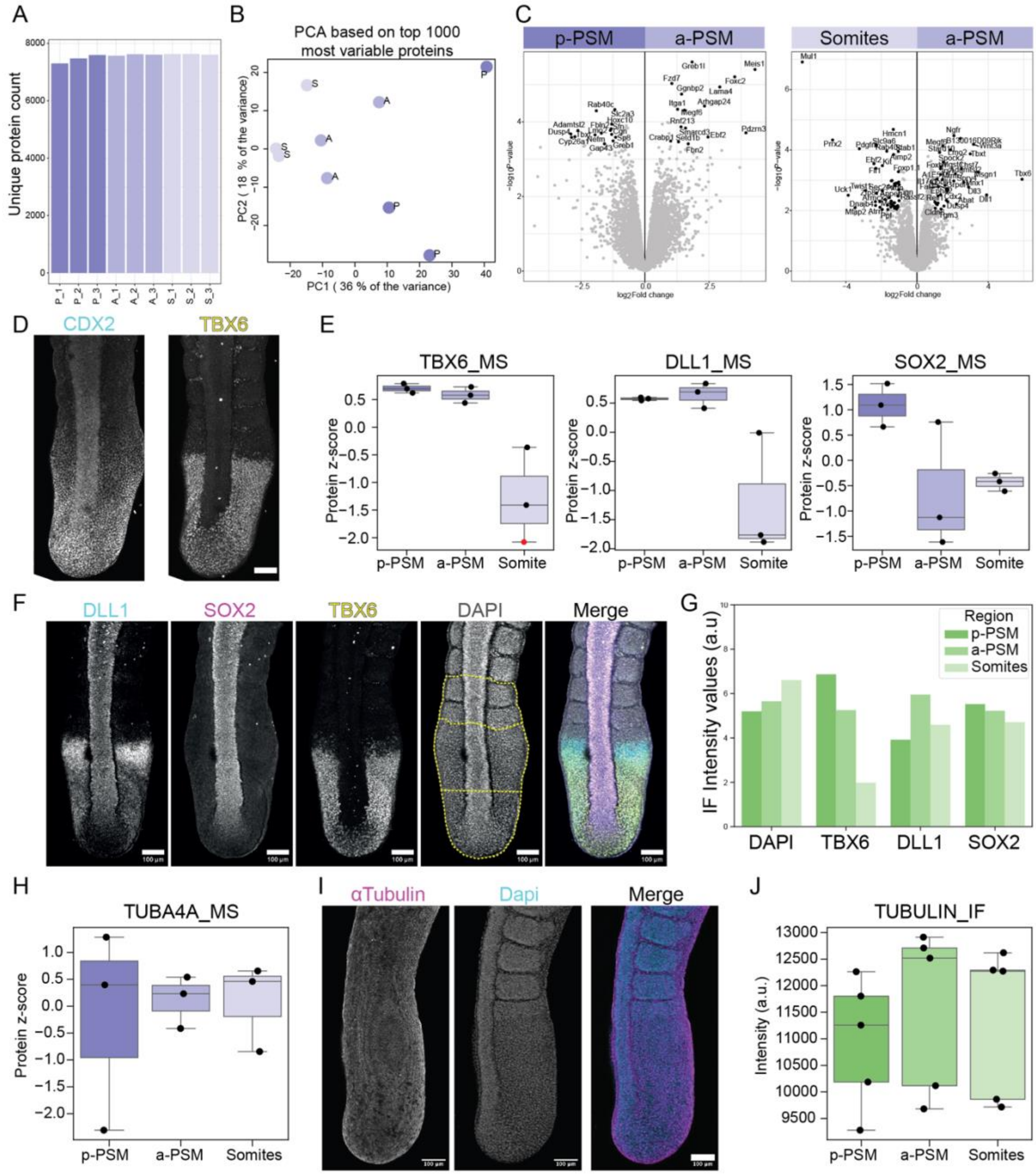


**Extended Data Figure 1, related to Figure 1. Analyses and validation of the proteomics data set.**

**A** Unique proteins measured in each sample replicate in the proteomics data set. P_1, A_1, S_1etc. : p-PSM, a-PSM, somite replicate 1, respectively. **B** PCA clustering of all sample replicates based on the top 1000 most variable proteins. **C** Volcano plot comparing the fold change from a-PSM to p-PSM (*left*) and a-PSM to Somite (*right*) in proteomics data set. **D**

Immunofluorescence of TBX6 and CDX2 (mesodermal markers) in intact uncut tails (corresponds to tail in Fig. 1B). **E** Mass spectrometry (MS) values corresponding to TBX6, DLL1, and SOX2 in p-PSM, a-PSM and Somite. Imputed value indicated with red dot. **F, G** Immunofluorescence (IF) of TBX6, DLL1, and SOX2 **F** and corresponding quantification of intensity in the p-PSM, a-PSM, Somites using a hand-drawn ROI around each region as shown in example yellow boxes in DAPI signal **F**. **H** MS values corresponding to Tubulin 4 alpha subunit. **I, J** Immunofluorescence of alpha-Tubulin in mouse embryo tail (**I**), and corresponding quantification for each region (**J**)**,** as done in **F**.

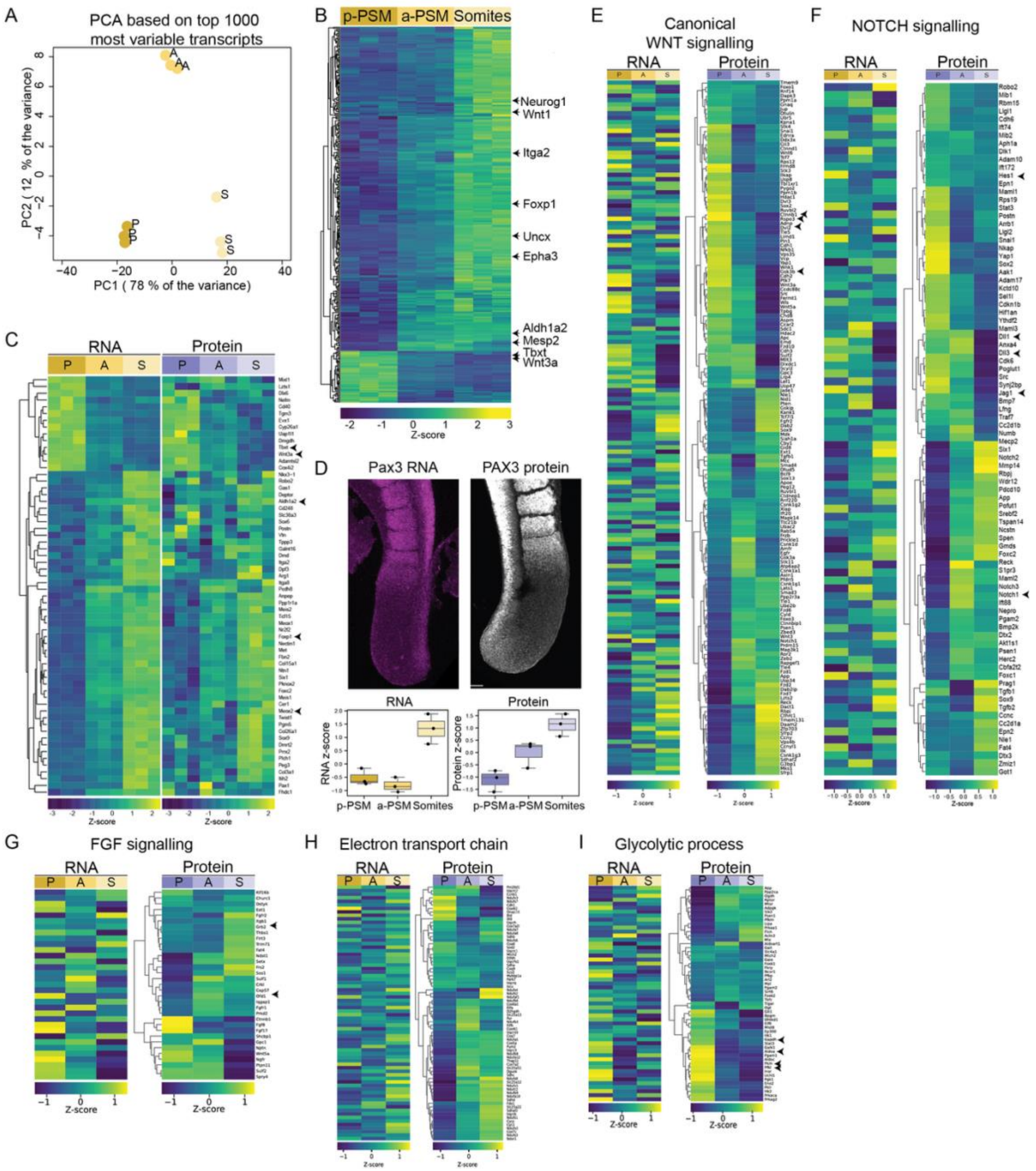


**Extended Data Figure 2, related to Figure 2. Spatial proteome and transcriptome patterns do not always overlap.**

**A** PCA clustering of all sample replicates based on the top 1000 most variable transcripts. **B** Clustermap of differentially expressed genes from bulk RNA sequencing. **C** Cluster heatmap of proteomics output for each transcript that is differentially expressed in the posterior (P), anterior (A) and somite (S) region. **D** *Top: Pax3* transcript (HCR, *left*) and protein staining (immunofluorescence, *right*, same as main Fig. 4J). *Bottom*: RNA and protein patterns for Pax3 from RNA-seq and proteomics data. Scale bar 100 mm. **E-I** Heatmaps of RNA-seq and proteomics datasets, ordered according to proteomics data, for the following designated GO terms: Canonical Wnt signalling (**E**, GO: 0060070), Notch Signalling (**F**, GO: 0007219), FGF (**G**, GO: 0008543), Electron transport chain (**H**, GO: 0022900) and Glycolytic process (**I**, GO: 0006096).

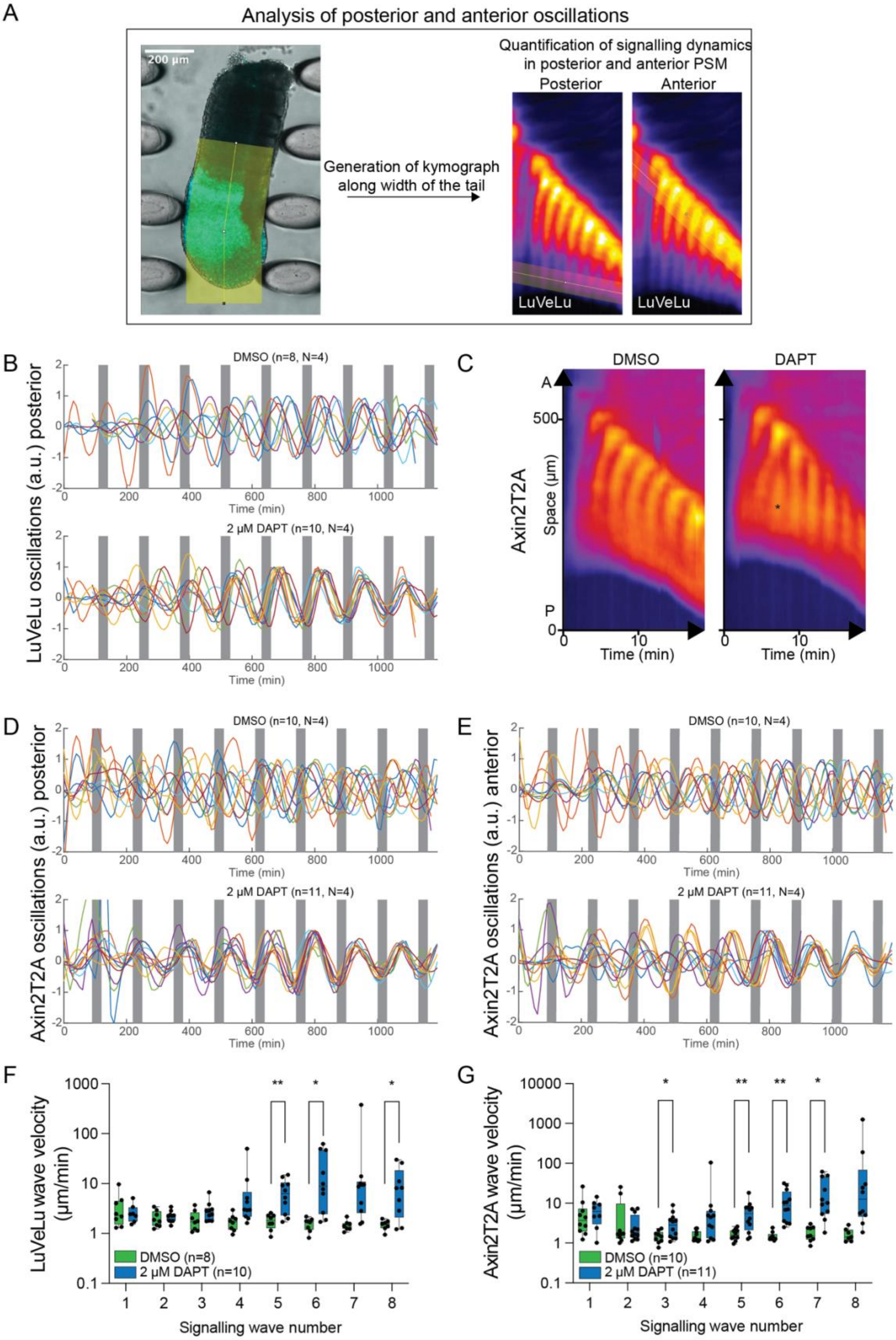
A
Analysis of posterior and anterior oscillations
200 µm
Generation of kymograph along width of the tail
Quantification of signalling dynamics in posterior and anterior PSM
Posterior
Anterior
LuVeLu
B
LuVeLu oscillations (a.u.) posterior
DMSO (n=8, N=4)
2 µM DAPT (n=10, N=4)
Time (min)
C
DMSO
DAPT
Axin2T2A
Space (µm)
A
P
500
Time (min)
D
Axin2T2A oscillations (a.u.) posterior
DMSO (n=10, N=4)
2 µM DAPT (n=11, N=4)
Time (min)
E
Axin2T2A oscillations (a.u.) anterior
DMSO (n=10, N=4)
2 µM DAPT (n=11, N=4)
Time (min)
F
LuVeLu wave velocity (µm/min)
DMSO (n=8)
2 µM DAPT (n=10)
Signalling wave number
G
Axin2T2A wave velocity (µm/min)
DMSO (n=10)
2 µM DAPT (n=11)
Signalling wave number

60

**Extended Data Figure 3, related to Figure 3. Entrainment of Axin2T2A mouse E10.5 embryonic tails upon DAPT treatment.**

**A** Representative image of oscillating LuVeLu kymograph obtained from manual drawing line scan in Fiji from posterior to anterior region of E10.5 mouse embryonic tails. Representative images of lines drawn in either the posterior or anterior region in the kymograph to obtain oscillatory behaviour in either region for analysis. **B** Detrended LuVeLu signal in anterior PSM (quantified from kymographs as described in **A**). Gray vertical bars in plots indicate the moment of medium / drug pulsing. **C** Representative kymographs for Axin2-T2A-Venus tails entrained with DMSO (*left*) or DAPT (*right*). **D, E** Detrended Axin2T2A signal in in the posterior (**D**) and anterior regions (**E**) (quantified from kymographs as described in **A**). Gray vertical bars in plots indicate the moment of medium / drug pulsing. **F, G** Quantification of wave speeds of LuVeLu oscillations (**F**) and Axin2-T2A (**G**). Absolute values were considered. (Unpaired T Test, * $p<0.05$, ** $p< 0.01$, rest non-significant)

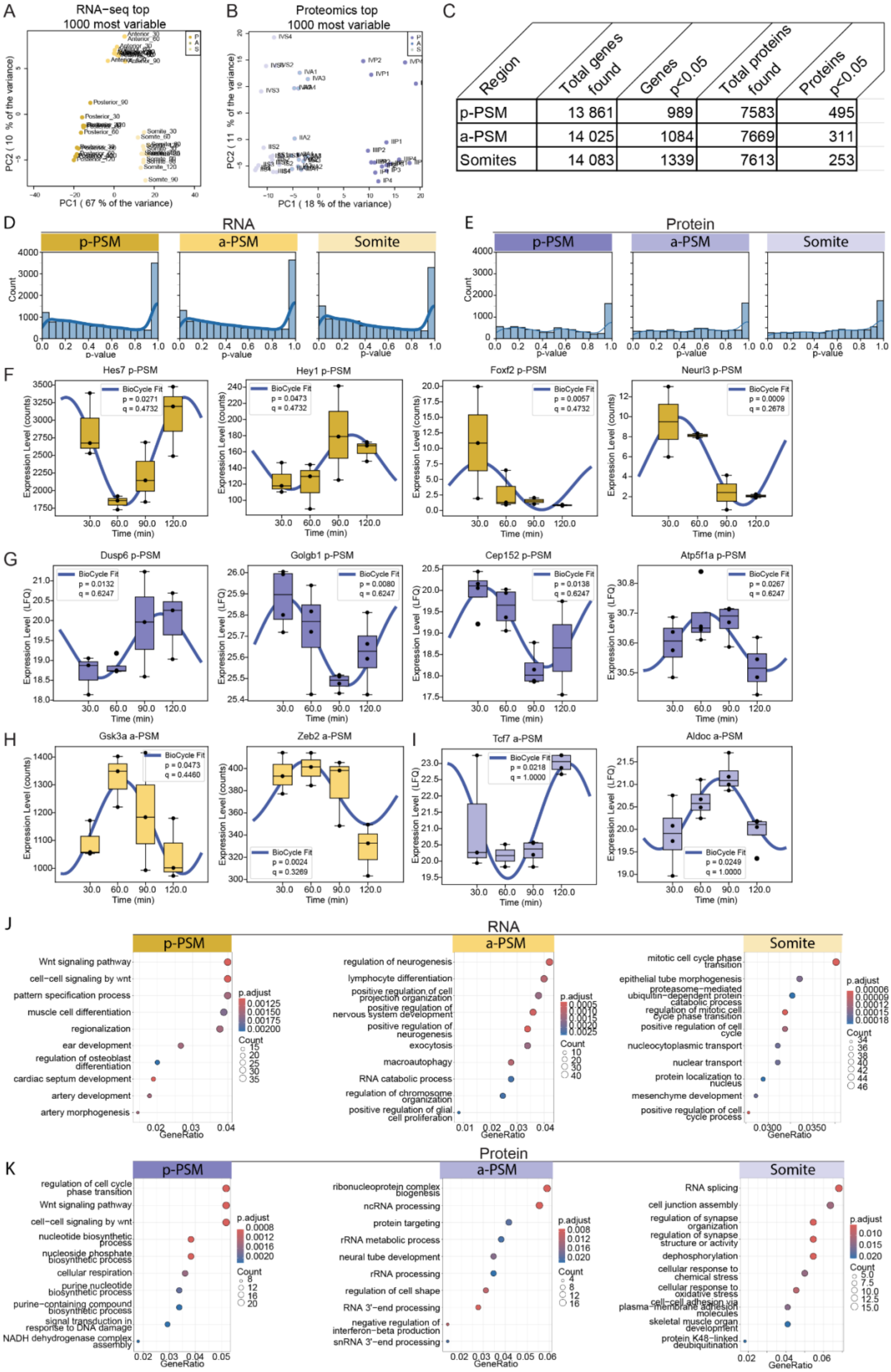


| Region | Total genes found | Genes p<0.05 | Total proteins found | Proteins p<0.05 |
|---|---|---|---|---|
| p-PSM | 13 861 | 989 | 7583 | 495 |
| a-PSM | 14 025 | 1084 | 7669 | 311 |
| Somites | 14 083 | 1339 | 7613 | 253 |

**Extended Data Figure 4, related to Figure 4. Identification of dynamic proteins and transcripts in the mouse embryo tail.**

**A, B** PCA clustering of all sample replicates for spatiotemporal RNA-seq (**A**) and proteomics (**B**) datasets based on the top 1000 most variable transcripts and proteins. **C** Summary table of total amount of transcripts and proteins and the significantly dynamic ones found after BIO-CYCLE analyses. **D, E** Distribution of obtained p-values after BIO-CYCLE analyses in the p-PSM, a-PSM and Somite region in the RNA-seq (**D**) datasets and proteomics (**E**). **F-I** Examples of predicted dynamic transcripts (**F**, **H**) and proteins (**G**, **I**): **F**, **G** Examples for p-PSM. Note the identification of known cyclic genes and new ones. **H, I** Further examples for a-PSM to Fig. 4. **J, K** Top 10 enriched GO terms within significantly dynamic transcripts (**J**) and proteins (**K**) in the indicated regions.

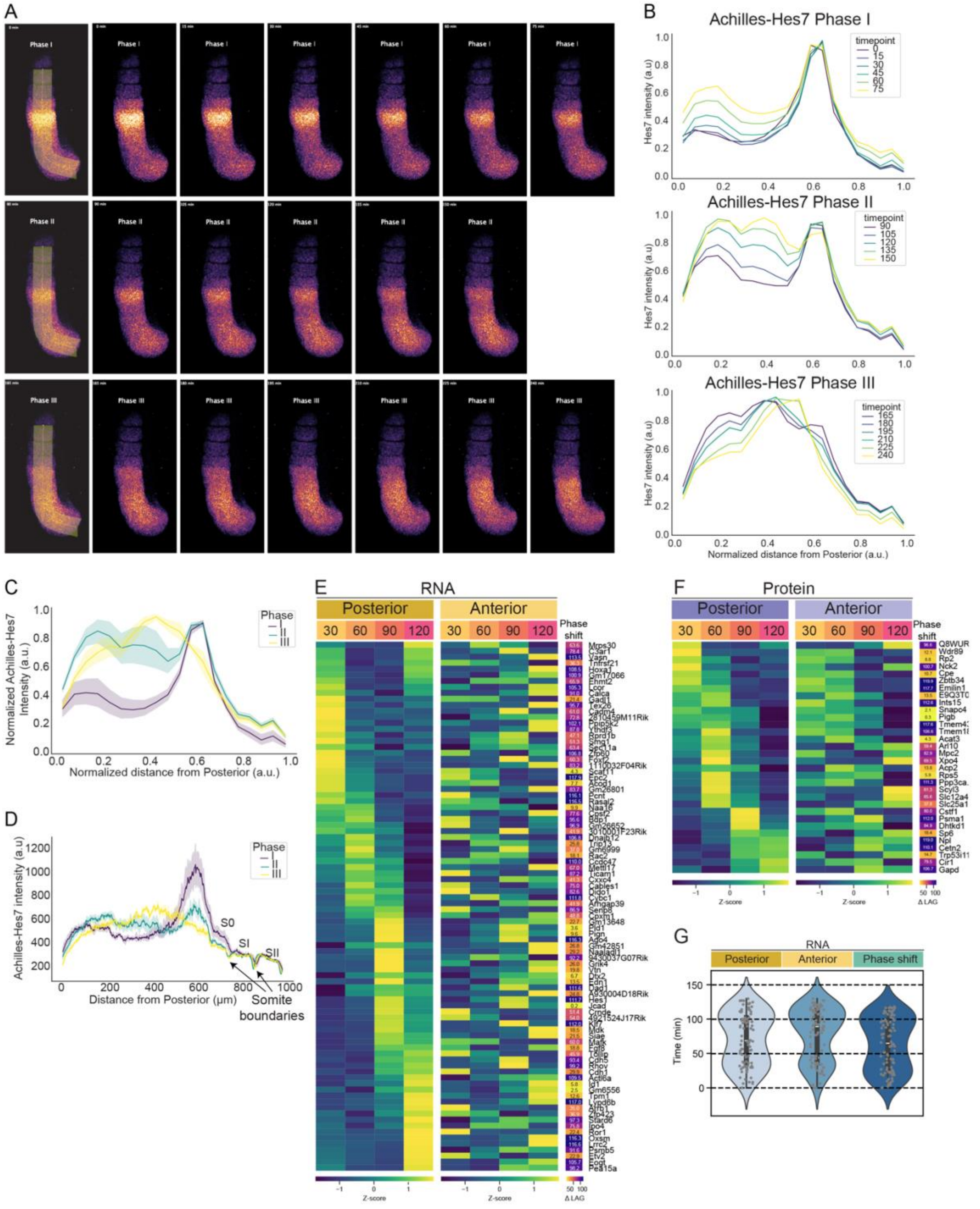


**Extended Data Figure 5, related to Figure 4. Hes7 oscillatory phases description.**

**A-D** Definition of Hes7 oscillation phases based on real-time imaging experiment of E10.5 mouse embryonic tail expressing Achilles-Hes7. **A** Snapshots of consecutive timepoints of one oscillation cycle were divided into three defined phases: Phase I: wave reaches the new anterior boundary next to S0; Phase II: decreasing intensity of Achilles-Hes7 at the new boundary and new wave starting at the posterior region; Phase III: new wave in the middle of the PSM. **B** Quantification of Achilles-Hes7 intensity along line indicated in **A** for each time point (color-coded), subdivided by phase. Intensity normalized to maximum per timepoint and space normalized between 0 (posterior end) and 1 (somite boundary). **C** Normalized average and SEM of data shown in **B**. **D** Raw Achilles-Hes7 intensity over distance from the posterior in the three Hes7-phases (average and SEM per phase shown). **E** Heatmap comparing significantly dynamic transcripts ($p<0.05$) found in both p-PSM and a-PSM and calculated phase-shift in min (phase-shift $_{\text{transcripts}}$ = lag time (a-PSM - p-PSM)). **F** Heatmap comparing significantly dynamic proteins ($p<0.05$) found in both p-PSM and a-PSM and calculated phase-shift in min (phase-shift $_{\text{proteins}}$ = lag time (a-PSM - p-PSM)). **G** Violin plot of lag time (min) for all significantly dynamic transcripts detected using BYO_CYCLE ($p<0.05$) in p-PSM and a-PSM and phase-shift (min) between the two.

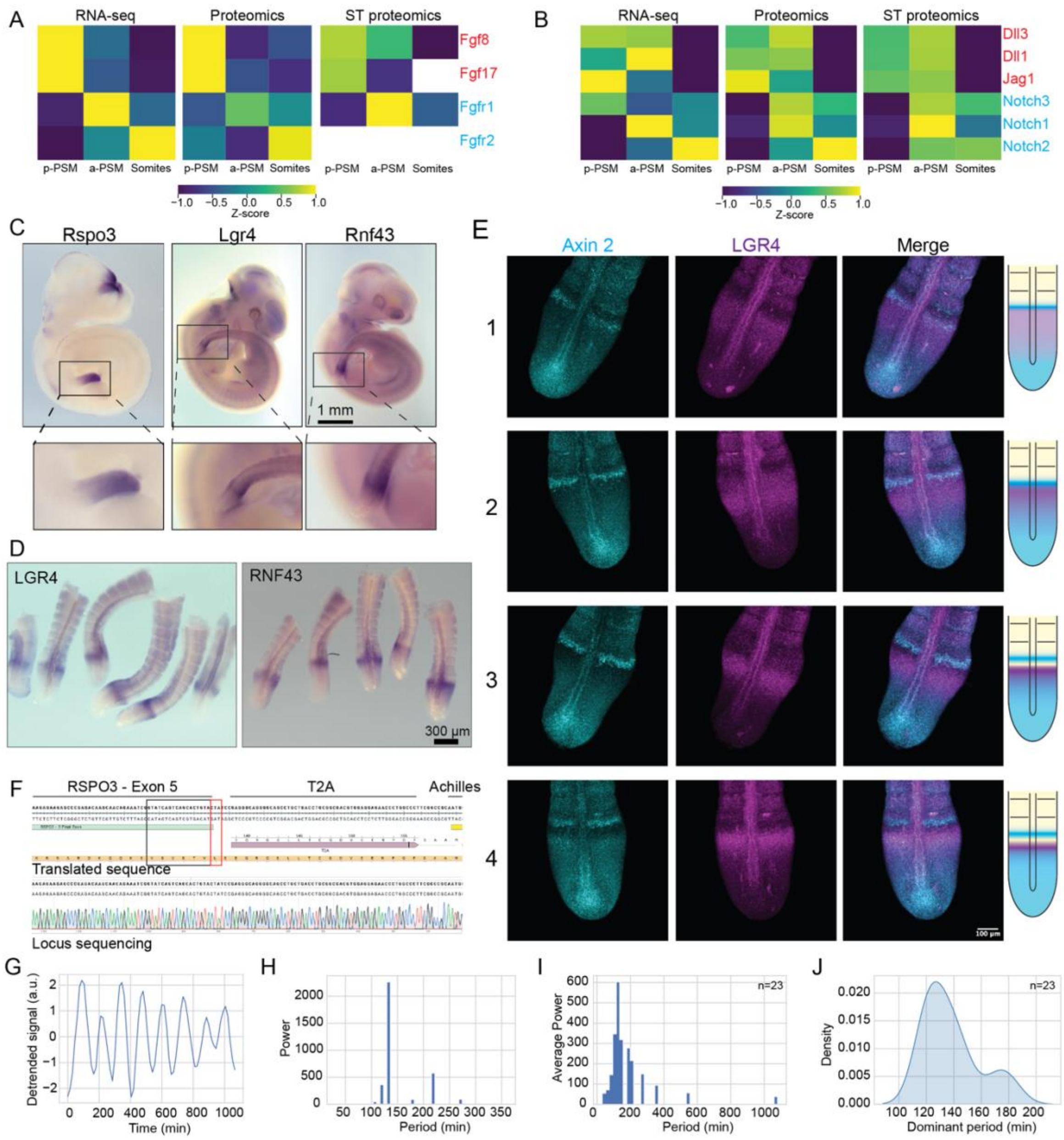


**Extended Data Figure 6, related to Figure 5. LGR4 and RSPO3 are antagonistically expressed.**

**A, B** Heatmap of selected components of FGF (**A**) and Notch (**B**) signalling pathways with annotated ligand (red) and receptors (cyan) found in the spatial RNA-seq and spatial and spatiotemporal (ST) proteomics datasets. **C, D** Representative images of *in situ* hybridizations of E10.5 mouse embryos for *Rspo3*, *Lgr4* and *Rnf43* (**C**) and dissected pooled tails for Lgr4 and Rnf43 (**D**). **E** Representative images of hybridization chain reactions (HCR) of E10.5 mouse embryonic tails for *Lgr4*, *Rspo3*, *Axin2* for four defined sequential *Axin2* wave oscillations pattern. Note the progressively brighter and more concentrated *Lgr4* stripe in anterior PSM corresponding to *Axin2* dynamics. Schematic summary of pattern for *Axin2* and *Lgr4* shown on the *right*. **F** Aligned Sanger sequencing results of genomic *Rspo3* locus edited with T2A-Achilles. Black box: last nucleotides of endogenous *Rspo3*. Due to nucleotide substitutions at the 3' end, there is an amino acid substitution from histidine to valine (red box). Translated sequence shown in yellow and locus sequencing is below. **G-J** Analysis of RSPO3-T2A-Achilles signal in gastruloids: **G, H** Representative detrended signal (**G**) and corresponding Fourier power spectrum (**H**) of RSPO3-T2A-Achilles signal in anterior PSM of one gastruloid. **I, J** Average Fourier power spectrum for all tracks analysed (**I**) and corresponding kernel density estimate (KDE) plot of dominant oscillation periods (**J**).

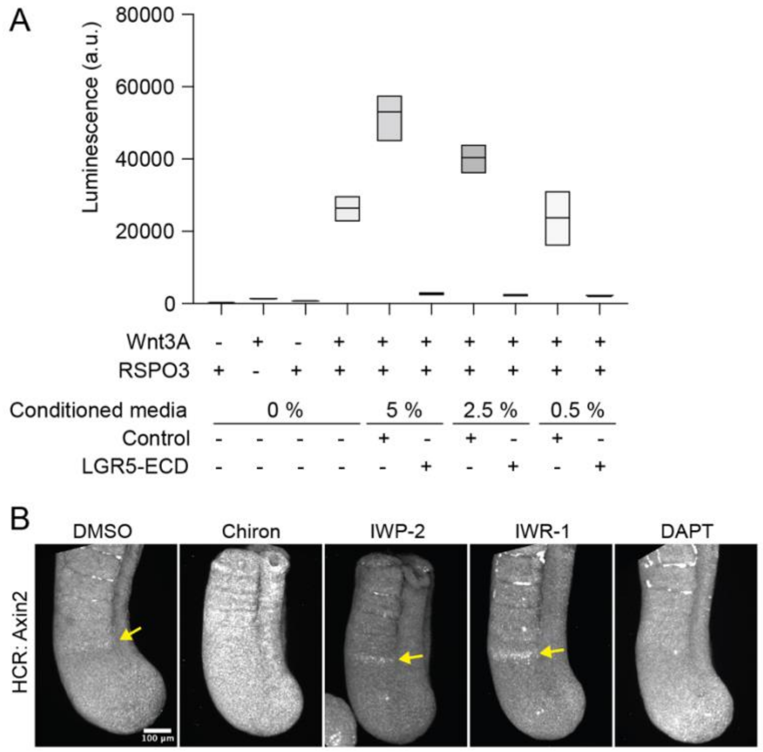


**Extended Data Figure 7, related to Figure 6. Effect of signalling perturbation on Wnt signalling and Axin2 expression.**

**A** Quantification of Wnt signalling activity (luminescence) in HEK293 cells expressing Super TOPFlash (STF) reporter in the presence of Wnt3A-conditioned medium and RSPO3. Note the expected induction in the presence of both Wnt3A and RSPO3. Control- and LGR5-ECD-conditioned media were added at the indicated concentrations. **B** Representative images of an *Axin2* HCR in E10.5 mouse embryo tails after 5 hours of *ex vivo* culture incubated with small molecule inhibitors targeting the Wnt and Notch pathways. DMSO (control), Chiron (10 µM, agonist of Wnt), IWP-2 (5 µM, Wnt inhibitor), IWR-1 (10 µM, Wnt inhibitor), DAPT (10 µM, Notch inhibitor) (N=4, $n_{DMSO}$=6, $n_{Chiron}$=6, $n_{IWP-2}$=5, $n_{IWR-1}$=6, $n_{DAPT}$=5). Yellow arrow: Axin2 stripe in forming somites.

### Supplementary Information

**Supplementary Data 1:** Heatmap of differentially expressed proteins

**Supplementary Data 2:** Heatmap of differentially expressed transcripts

**Supplementary Video 1.** Embryonic tails expressing LuVeLu reporter upon entrainment with DMSO control or DAPT, related to Fig. 3.

**Supplementary Video 2.** Mouse embryonic tail expressing Achilles-Hes7 over one oscillation period, related to Fig. 4.

**Supplementary Video 3.** Real-time imaging of mouse gastruloids expressing RSPO3-T2A-Achilles reporter, related to Fig. 5.

**Supplementary Video 4.** Spread-out cultures of mouse embryonic tails expressing Axin2-T2A-Venus treated with control (*left*), RSPO3 (*middle*) or LGR5-ECD (*right*), related to Fig. 6.
