## Supplemental Information for "Spatiotemporal proteomics reveals dynamic antagonistic gradients shaping signalling waves"

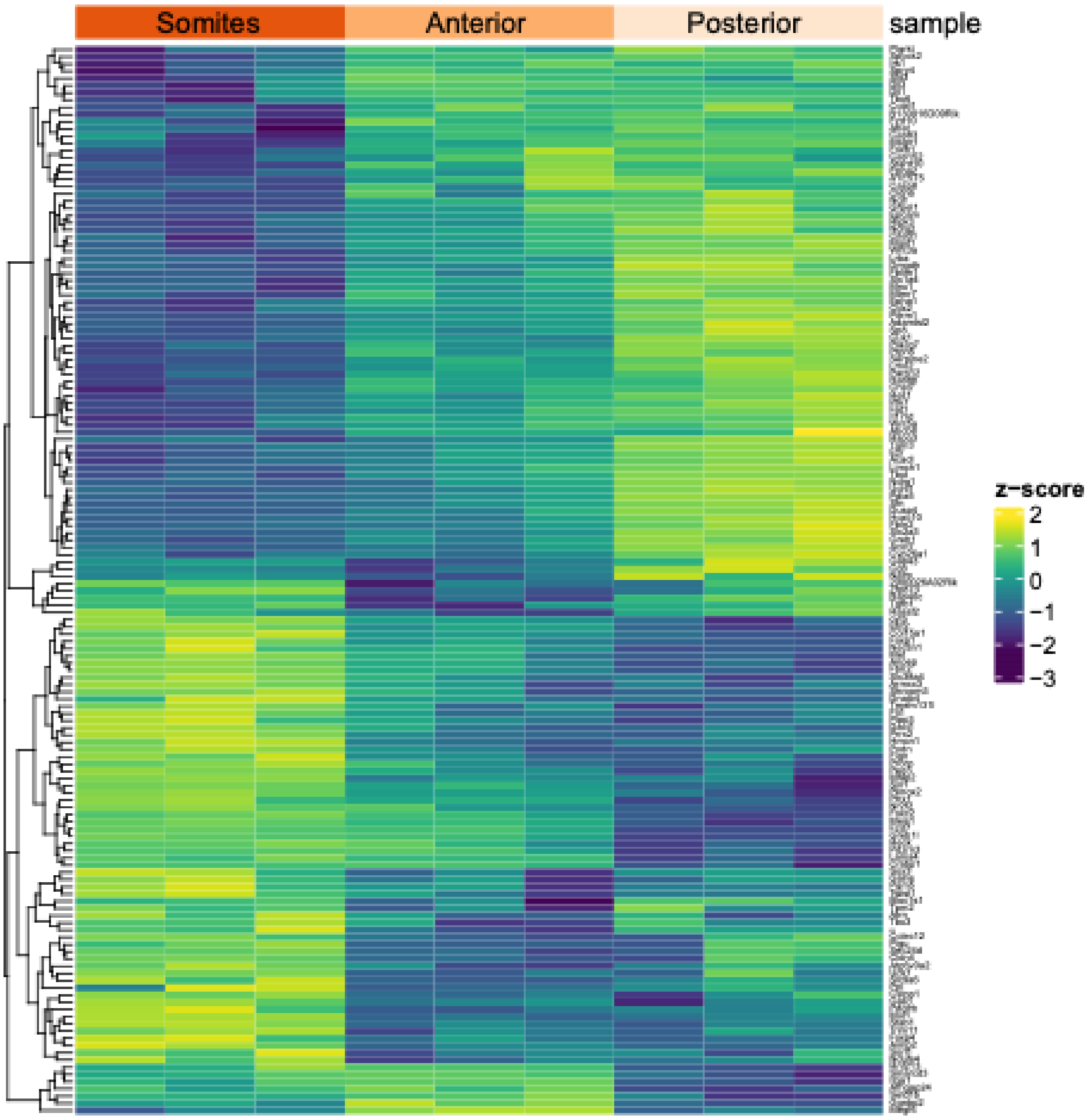


**Supplementary Figure 1 –** Heatmap of total differentially expressed proteins

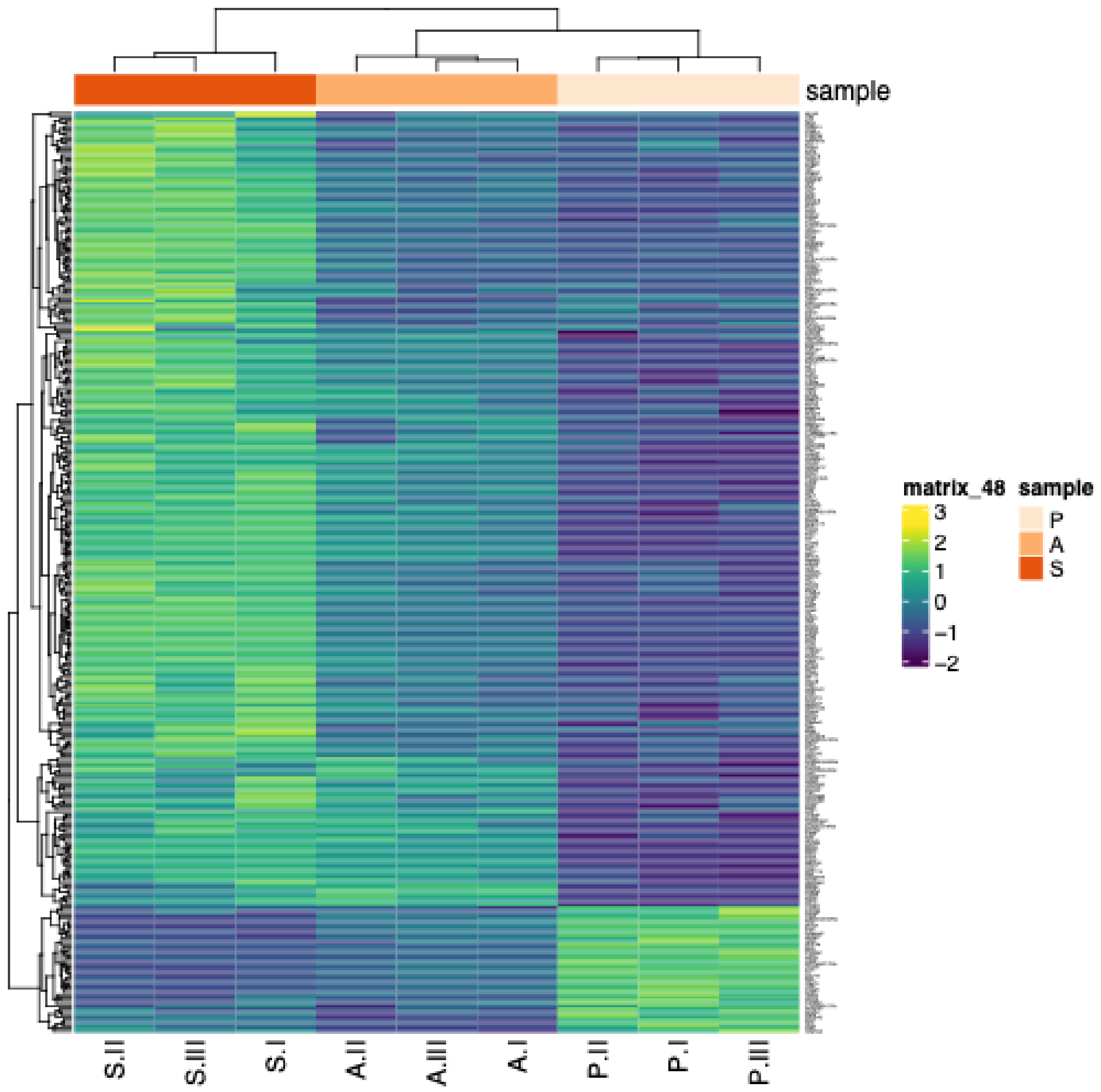


**Supplementary Figure 2** – Heatmap of total differentially expressed transcripts
